# Tumor Cells Enriched for Interferon and Inflammatory Programs Pre-Exist in High Grade Serous Ovarian Cancer and are Proportionately Significantly Increased Post Chemotherapy

**DOI:** 10.1101/2025.06.24.661404

**Authors:** Boris J.N. Winterhoff, Nomeda Girnius, Muyi Liu, Jaeyoung Kang, Mihir Shetty, M. Angie Martinez-Gakidis, Weiyi Xu, Mayura Thomas, Linna Lahmadi, Aylin Z. Henstridge, Pilar Baldominos, Shobhana Talukdaer, Zenas Chang, Jordan Mattson, Morgan Gruner, Sally A. Mullany, Peter Argenta, Gottfried E. Konecny, Dennis J. Slamon, Ronny Drapkin, Andrew C. Nelson, Timothy K. Starr, Joan S. Brugge

**Author notes:** These authors contributed equally (Boris J.N. Winterhoff, Nomeda Girnius, Muyi Liu).

## Abstract

Drug-tolerant, high-grade serous ovarian cancer (HGSOC) cells that persist after first-line chemotherapy and subsequently relapse often retain sensitivity to secondary treatment, suggesting a therapeutic window before stable chemoresistance emerges. We performed single-cell RNA sequencing (scRNA-seq) on seven matched pairs of tumors, collected pre- and post-chemotherapy, to define vulnerabilities in these reversibly-resistant cells. Treatment induced a marked enrichment of tumor and stromal cell populations expressing correlated interferon (IFN) and inflammatory (IFM) gene signatures, with a concurrent depletion of proliferation-related and MYC-associated states in the tumor cells. Cross-cohort single cell sequencing analysis of >130 treatment-naive tumors revealed heterogeneity in the abundance of IFN/IFM-expressing cells. Multiplex immunofluorescence imaging of IFN-stimulated gene (ISG) products confirmed the presence of spatially clustered ISG-positive tumor cells in all cases, as well as in serous tubal intraepithelial carcinomas (STIC lesions), the presumptive HGSOC precursors. ISG expression correlated strongly with ORF1P, a protein encoded by the endogenous retrotransposon LINE1. These data suggest that early oncogenic events drive LINE1 derepression and innate immune activation, establishing an IFN-rich transcriptional state that persists in tumor subpopulations and is strongly enhanced by chemotherapy.

## Main

While 80-85% of high grade serous ovarian cancers (HGSOC) undergo dramatic regression with platinum/taxane-based chemotherapies, the majority of tumors relapse within 18 months and are no longer curable^1–3^. Recurrent tumors initially display sensitivity to repeated courses of chemotherapy, but continued relapses become increasingly resistant to treatment. Improving chemotherapy outcomes is critically dependent on characterizing drug-tolerant cells to understand their heterogeneity and identify cellular programs key to development of resistance.

The genetic and phenotypic states of treatment-naive HGSOC tumors have to date been defined through bulk genomic, transcriptomic and proteomic analyses. HGSOC tumors are characterized by extensive copy number alterations, gene breakage and chromosomal rearrangements in the background of almost universal TP53 mutations^4–6^. While genes and proteins or multi-gene/protein signatures associated with outcome to chemotherapy have been identified by these studies (e.g. EMT, allograft rejection, AP1 transcription factors and IFN responses^7–12^), it is difficult to correlate most of these signatures with specific tumor-intrinsic programs because stroma cells are included in the bulk analyses and these chemotherapy-enriched genes/signatures can be expressed in many cell types. This is one of the reasons bulk RNA-based molecular classifiers (e.g. ^4,13–15^) have been difficult to translate into clinical use.

Single-cell RNA studies provide a more precise analysis of the cell types that comprise pre-treatment tumors as well as transcriptional states associated with these cell types^16–23^. Negative matrix factorization (NMF) and differential gene expression (DGE) analyses of cellular programs across patients has consistently defined a large set of modules with coherently co-varying gene expression which can be linked to specific cellular functions including cell cycle regulation, inflammation, stress response, antigen presentation, interferon response, and epithelial-mesenchymal transition^16,17,20^. Phenotypic variation is associated with different genomic subtypes of HGSOC (e.g. inflammatory programs enriched in tumors with homologous repair deficiency) as well as stromal states specific to anatomical sites^20^. In addition, whole genome doubling (WGD) is an ongoing mutation process that promotes cell diversity and affects the phenotypic state of tumor cells and the microenvironment^24^. Multiple studies defined specific subtypes of immune and stromal cell populations associated with tumors as well as intrinsic tumor cells programs that are enriched in neighborhoods of infiltrating T cells or immune cell-free neighborhoods^17,18,20,25,26^.

Cellular programs associated with platinum resistance in vitro include: DNA damage repair, drug detoxification, epithelial-mesenchymal transition, calcium binding proteins, chaperones, extracellular matrix, autophagy, metabolic enzymes, transcription factor, growth factor signal transduction among others (reviews^27,28^, and^29–32^). The extent to which each program is involved in the early development of drug tolerance has not been established. Only a few studies have used scRNAseq to analyze post-treatment samples from HGSOC prior to first relapse to identify intrinsic cellular programs of tumor cells that survive initial chemotherapy^33–36^.

In this study, we carried out scRNAseq and spatial imaging of tumors before and after chemotherapy exposure to focus on tumor-intrinsic cellular programs that are enriched in tumor cells which survive chemotherapy. Our data reveal an enrichment of canonical interferon program genes which show correlated expression with multiple other inflammation-associated gene programs. Tumor cells expressing these signatures are present in pre-treatment tumors across all metastatic sites with distinct proportional representations and are also present in precursor STIC lesions. The highly correlated expression of ISGs and the LINE1 protein ORF1P suggest that derepression of repeat elements is involved in triggering the IFN/IFM response.

## Results

### ScRNA-seq reveals enrichment of inflammatory programs in post-treatment tumors

To evaluate cancer cell-intrinsic programs that are altered after chemotherapy, patient-matched biopsies were collected from seven newly diagnosed, treatment-naive HGSOC patients during laparoscopic surgery (pre-treatment) and again during interval debulking surgery after neoadjuvant chemotherapy (NACT) (post-treatment). Tissue from each biopsy was either formalin-fixed for histological assessment and multiplex immunofluorescence imaging or dissociated to single cells and analyzed by droplet-based 10X Genomics scRNA-seq (Figure 1a). Disease progression and chemotherapy regimens varied considerably among the seven patients (Figure 1b and Extended Table 1).

**Figure 1:**
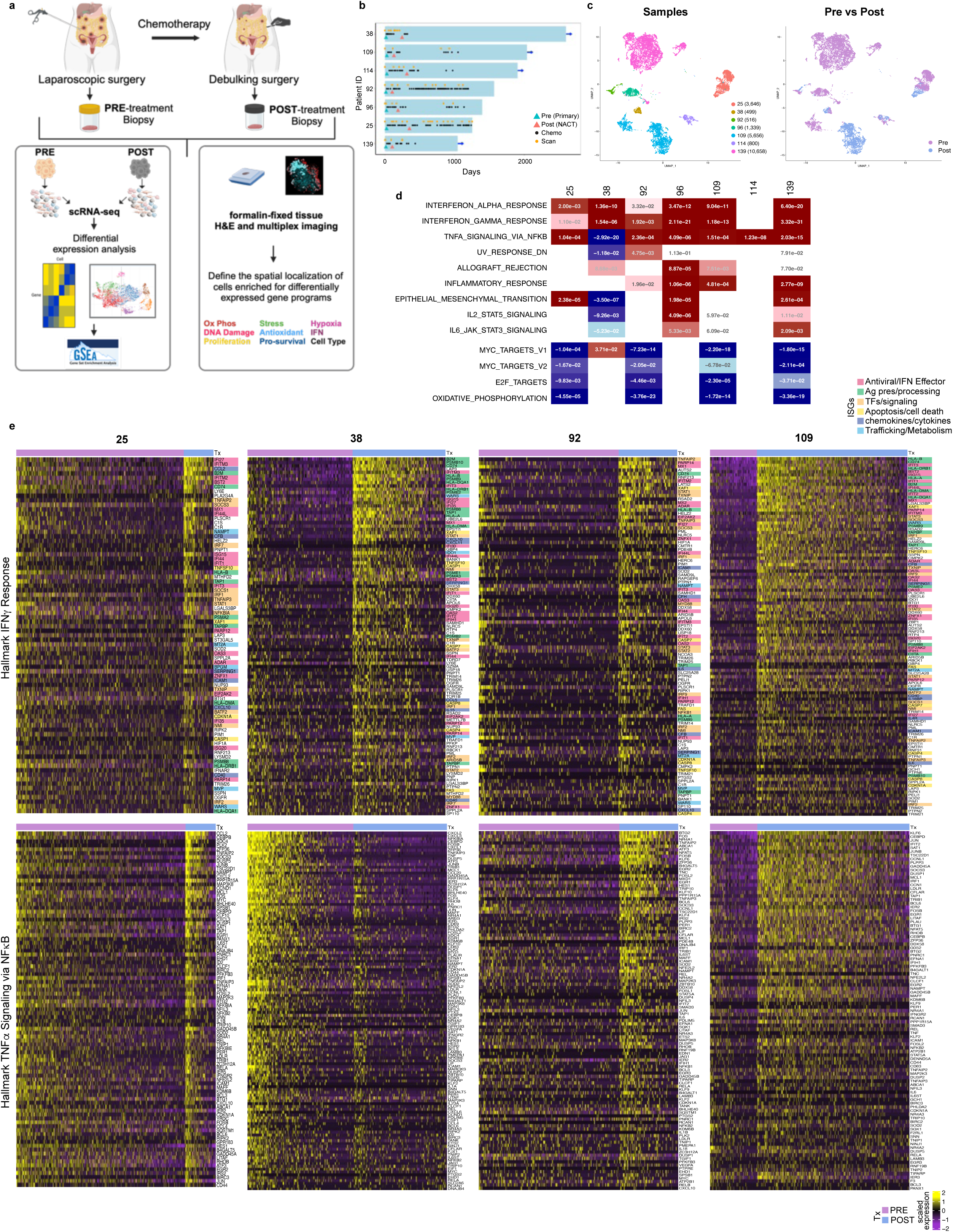
Analysis of differentially expressed genes in pre- and post-treatment HGSOC tumors. a. Schematic diagram of the experimental design. b. Overall survivor swimmers plot. Blue arrows indicate the patient is still alive. Green triangles = pre-NACT sample acquisition, Red triangles = post-NACT sample acquisition. Black dots = chemotherapy administration. Orange dots = diagnostic scans. c. Uniform manifold approximation and projection (UMAP) plots of single epithelial cells from all seven paired tumor samples color-coded by patient number (left panel) or by pre- vs post-NACT collection time (right panel). d. Table of adjusted p-values for Hallmark gene sets showing significant differential expression in post-treatment cells compared to pre-treatment cells for each tumor pair (red = up-regulation, blue = down-regulation). e. Heatmaps showing the level of expression of differentially expressed Hallmark IFNγ and ‘TNFα signaling via NFκB signature’ genes in the four paired samples with high post-treatment cell counts (patients 25, 38, 92 and 109). Cells were separated into pre- and post-treatment bins and rank ordered by their IFNγ and TNFα signature scores (left to right, decreasing order) (purple bar = pre; blue bar = post). Genes were rank-ordered in the maps based on the cluster average fold changes between post- and pre-treated tumors. Color coding represents the different classes of ISGs (see legend).

**Extended Table 1.** Patient and sample clinical data. ^a^Anatomical site: ome = omentum, ova = ovary, per = peritoneal cavity, abd = abdomen. ^b^Ptx response = response to platinum chemotherapy, defined as sensitive if there is no relapse within six months of chemotherapy. ^3^RD = Residual disease, based on surgeon’s assessment after debulking surgery. R0 = no visible disease, R1 = residual disease < 1 cm in size. ^d^CRS = Chemotherapy response score. Pathologist scores by comparing pre- and post-NACT pathology: 1 = no or minimal tumor response, 2 = partial tumor response, 3 = complete or near-complete response (PMID: 31482065). ^e^Stage = FIGO staging. ^f^Days = days to relapse after initial diagnosis or days to last follow-up, if patient has not relapsed (indicated with *)

| ID | Pre site<sup>a</sup> | Post site<sup>a</sup> | Ptx response<sup>b</sup> | RD<sup>c</sup> | CRS<sup>d</sup> | Stage<sup>e</sup> | Relapse | Days<sup>f</sup> |
| --- | --- | --- | --- | --- | --- | --- | --- | --- |
| 25 | ome | ome | resistant | R0 | 1 | IVB | Yes | 556 |
| 38 | ome | ome | sensitive | R0 | 3 | IVB | No | 2598* |
| 92 | ome | ova | resistant | R1 | 2 | IV | Yes | 330 |
| 96 | per | ome | sensitive | R0 | 3 | IIIB | Yes | 867 |
| 109 | ome | ova | sensitive | R0 | 3 | IV | Yes | 806 |
| 114 | ome | ome | sensitive | R0 | 1 | IV | No | 1903* |
| 139 | per | abd | sensitive | R0 | 1 | IVB | Yes | 1121 |

Epithelial cells were identified in the scRNA-seq dataset and clustered to evaluate transcriptional relatedness. As previously shown in other studies, each tumor formed distinct transcription clusters^16–18,20^ (Figure 1c). To evaluate gene expression differences resulting from chemotherapy treatment, we compared each pair of pre- and post-treatment samples from individual patients separately and identified differentially expressed genes (DEGs) (Supplementary Table 1). To identify gene signatures associated with chemotherapy, we performed gene set enrichment analysis (GSEA) on these gene lists. In six of the seven pairs, there was significant enrichment of Hallmark IFNγ and Inflammatory response signatures as well as TNFα signaling via NFκB (TNFA/NFKB) signature in the post-treatment samples (Figure 1d, and Supplementary Table 2). Unlike other cases, patient tumor 38 showed strong enrichment in the TNFA/NFKB and other inflammatory signatures in the pretreatment biopsy (Figure 1d). This is particularly interesting because this patient had severe peritonitis, which could provoke the high inflammatory signature. Interestingly her neoadjuvant course was complicated by severe peritonitis and immunofluorescence analysis revealed numerous small patches of CD11b+ immune cell clusters throughout her pretreatment sample, consistent with a highly inflamed tumor prior to chemotherapy (Extended data figure 1a).

To evaluate the most commonly enriched programs in more detail, we generated heatmaps of the enriched genes from the IFNγ and TNFA/NFKB signatures (Figure 1e and Extended Figure 1b). We tracked Hallmark IFNγ, rather than IFNα, signature genes because there is significant overlap in the genes from these two signatures and the IFNγ signature is more inclusive of genes commonly associated with either IFNγ or IFNα response (Extended data figure 1c).

Heatmaps of the enriched IFNγ and TNFA/NFKB signatures clearly demonstrated that there is coordinated expression of a large number of bona fide canonical IFN and TNFA/NFKB signature genes in a majority of post-treatment cells. The inflammatory gene signature heatmap in Figure 1e also validated the high expression of inflammatory genes in pre-treatment tumor 38. The heatmaps also show that a variable number of cells from the pre-treatment biopsies also displayed coordinated expression of multiple IFNγ and TNFA/NFKB signature genes, suggesting that cells enriched for these signatures pre-exist in pretreatment tumors (Figure 1e, Extended Figure 1b).

The most commonly enriched IFNγ, IFNα, and TNFα signature genes across all the samples are shown in Supplementary Table 4. While a high percentage of canonical IFN pathway genes are recurrently enriched among the IFNγ signatures, a large percentage of the recurrent TNFA/NFKB signature genes are IEGs which are commonly induced after stimulation of many receptors, but can also be induced during tissue dissociation for scRNAseq analysis^37,38^.

Three other inflammation-related signatures were also enriched in a subset of the post-treatment samples (Inflammatory response, IL2-STAT5 signaling, and IL6-JAK-STAT3) (Figure 1d). The DEGs in the Allograft Rejection signature were predominantly IFN-induced genes involved in antigen processing and presentation (HLA class I and II genes, CD74, B2M, TAP1/2) (Supplementary table 2). In patient 114 tumors, there was a comparable proportion of cells enriched for IFN signatures in pre- and post-treatment tumors, yet the TNFA/NFKB signature was enriched post-treatment (Extended Figure 1b).

### IFNγ signature is positively correlated with inflammatory signatures and negatively correlated with proliferation-related signatures in single cells

The evidence that both the IFN and TNFA/NFKB signatures were induced in a large proportion of post-treatment tumor cells suggests that these signatures may be strongly correlated in individual cells. We analyzed the correlation between GSEA scores for the IFNγ hallmark signature and 50 Hallmark signatures, a PCNA proliferation signature and a GO integrated stress signature in every pre- and post-treatment cell (Figure 2a). Multiple signatures associated with inflammation were correlated with the IFNγ hallmark signature (Pearson correlation, r > 0.25), including Inflammatory response, IFNα response, Complement, IL6-JAK-STAT3_signaling, TNFα signaling via NFκB, and IL2-STAT5 signaling (Figure 2b). The most strongly anti-correlated signatures consisted of proliferation-associated signatures including MYC targets, E2F targets, PCNA metagene, G2M checkpoint and Mitotic spindle (Figure 2c), suggesting that cells enriched for IFN and inflammation-associated signatures are more quiescent. Oxidative phosphorylation and DNA repair signatures were also anti-correlated with these programs in many samples. The lack of a correlation with any DNA damage signatures was supported by the failure to detect the DNA damage reporter γH2AX in cells expressing the IFN-induce protein IFIT1 *in situ* (Supplemental data Figure 1a-b), and BAF1+ micronuclei, indicative of chromosome instability or DNA damage^39^, were not associated with IFIT1 expression (Supplemental data Figure 1c). Interestingly, EMT and Hypoxia programs were only correlated in a subset of samples (EMT: 25, 109, 114, 139; hypoxia: 25, 38, 139). The evidence that multiple different signatures which are regulated by distinct transcription factors are coordinately expressed in the same cells suggests that there may be widespread epigenetic alterations that open chromatin to a large number of transcription factors.

**Figure 2:**
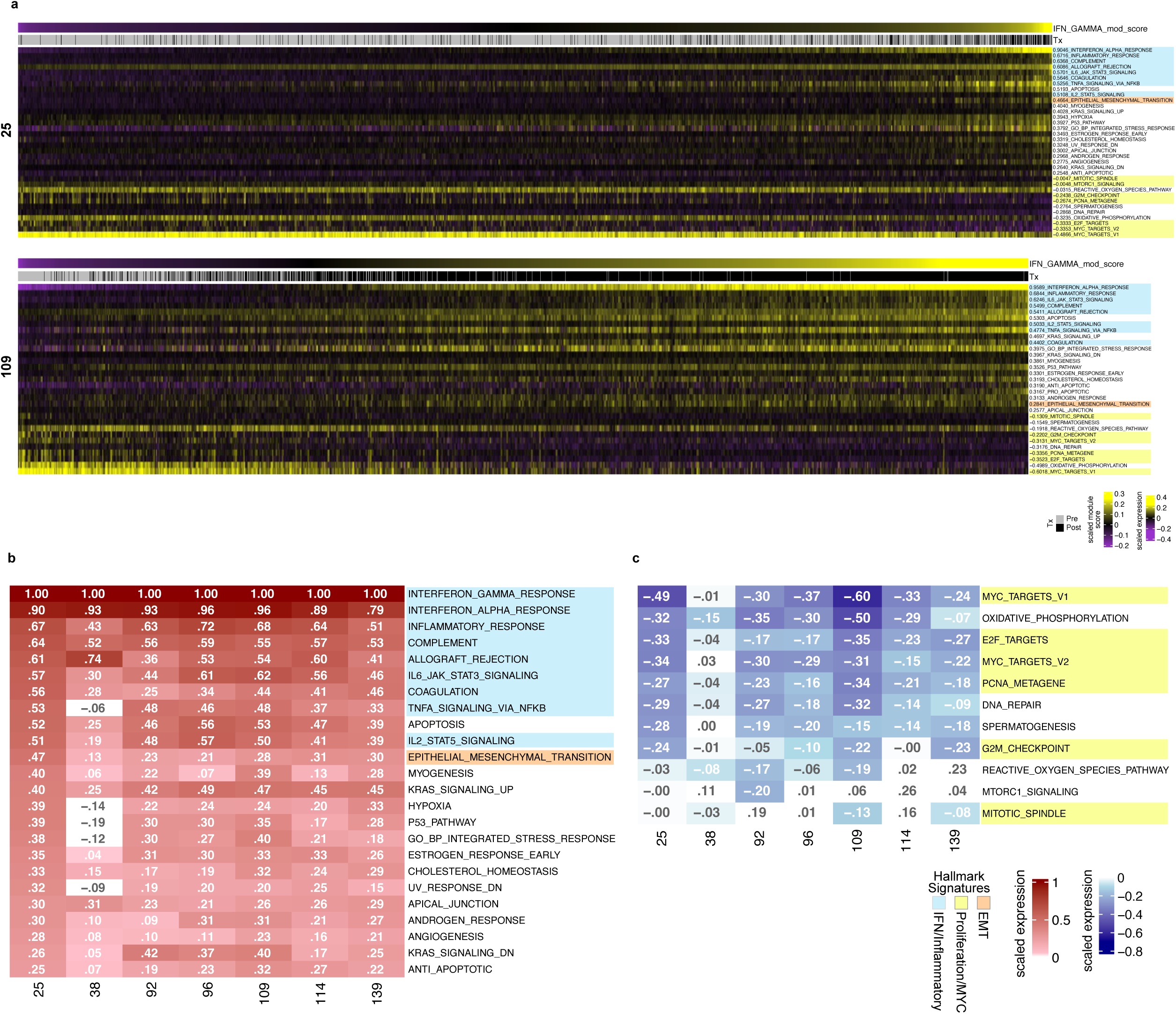
Signatures that correlate with IFNγ. a. Heatmap of GSEA signature scores of tumor cells from two representative pre- and post-tumor pairs (upper 25; lower 109). Cells are rank ordered based on their IFN*γ* signature scores (low on the left side, high on the right side) and the scores for each signature that was correlated or anti-correlated with IFNγ are shown for each cell. Positively correlated signatures (Pearson correlation, r > 0.25) and all negatively correlated signatures are shown. Heatmap colored tables of Pearson correlation values (IFNγ vs each signature) for each patient for positively correlated signatures (b) and negatively correlated signatures (c). IFN/IFM signature pathways are highlighted in blue, while proliferation-related signatures are highlighted in yellow. EMT is highlighted in orange.

To test whether or not these correlations were driven by common genes within the two signatures being compared, we calculated the overlap of genes between the various gene sets (Supplemental data Figure 1d). While there are a few pairs of signatures with some overlapping genes, the overlap between most others pairs of signatures is less than 5% suggesting overlap of common genes in the gene sets does not significantly contribute to the correlations, other than the predicted IFNα and IFNγ overlap.

### Subclusters of pre- and post-treatment cancer cells differ in their levels of IFN/IFM and proliferation-related signaling

We performed unsupervised cluster analysis of the tumor epithelial cells to provide a more detailed analysis of the subsets of cells that express IFN/IFM and proliferation signatures within each patient. Our analyses identified between 4-8 cancer cell clusters per patient, with individual clusters generally containing cells almost exclusively from either the pre- or the post-therapy samples (Figure 3a and Extended data figure 2a). In 5 of 7 patients, we detected at least two distinct post-treatment cancer cell clusters suggesting heterogeneity in cell populations that survive chemotherapy (Figure 3b and Extended data figure 2b). The violin plots show the IFNγ and PCNA module scores for each cluster. For this analysis, we used the IFNγ Hallmark gene set as a proxy for the IFN/IFM signatures and the PCNA metagene geneset as a proxy for proliferation-related signatures due to their high correlations with these signatures (Figure 2b).

**Figure 3:**
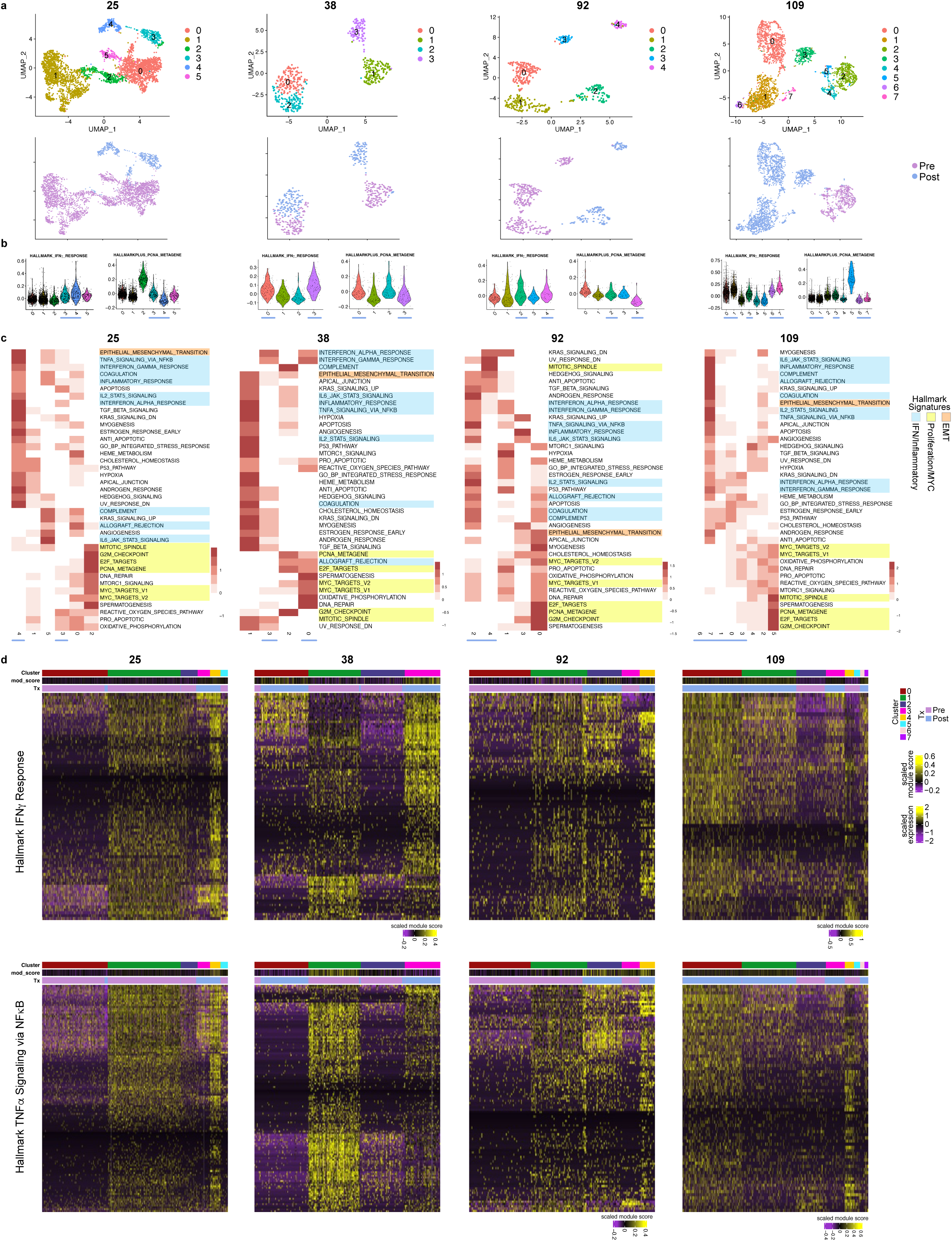
Analysis of cancer cell clusters in each pre-post tumor pair. a. UMAP plots of cancer cell clusters in four pre-post tumor pairs. Colors denote the cluster numbers (upper panels) or the pre-, post-treatment status of the cells (lower panels). b. Violin plots of the IFN*γ* and PCNA signature scores for each cluster. Blue bars are placed under clusters where the majority of cells are post-therapy. c. Heatmaps showing the signature scores for the pseudo-bulk average gene expression of each cluster for 39 gene sets. The 39 gene sets shown all had differential expression between the clusters and enrichment of canonical genes in their respective pathways (see Methods). Gene sets are colored by functional category (blue: IFN/IFM, yellow: proliferation-related, and orange: EMT). d. Heatmaps showing the expression of the Hallmark IFNγ response or TNFA/NFKB signature genes in each cluster. Cells in columns are arranged by cluster and genes in rows are clustered by level of expression. Scaled module scores for each cluster are shown in the top middle bar. The scale for each heatmap module score is as indicated on the right except for those cases where a different scale was required. These scale bars are underneath the heatmaps.

Consistent with the known sensitivity of proliferating cells to chemotherapy, the post-treatment clusters displayed low PCNA scores, with the exception of sample 38 where one post-treatment cluster contained contain cells with PCNA scores above the median. In addition, the clustering analysis demonstrated that the post-treatment clusters which had the most elevated IFNγ signature score had the lowest PCNA proliferation scores (Figure 3b,c and Extended data figure 2b,c) (patient 25 cluster 4 vs 3, patient 38 cluster 3 vs 0, patient 92 cluster 4 vs 2, and patient 109 cluster 7 vs 6), suggesting a lower proliferation rate and thus reduced sensitivity to chemotherapy.

To compare the levels of enrichment of gene signatures in cancer cell clusters within each patient, we performed GSEA using 50 Hallmark gene sets plus an additional eight curated datasets (Supplementary Table 5) (Figure 3c, Extended Figure 2c). Generally, one cluster from each pre- and post-tumor sample was enriched for the set of IFN-IFM-correlated signatures that were defined in the analysis in Fig. 2. The enrichment of these signatures is greater in the post-treatment clusters as predicted (Figure 3c). The heatmaps of IFNγ and TNFA/NFKB genes showed that these clusters displayed a highly coordinated pattern of expression of most genes in the IFN and TNFA/NFKB signatures (Figure 3d and Extended data figure 2d). It is noteworthy that a similar set of IFN- and TNFA/NFKB-signature genes are expressed in pre- and post-treatment clusters. For example, in patient 25 pre-treatment cluster 1 expressed a similar pattern of IFN and TNFA/NFKB genes pattern of expression as in post-treatment cluster 4 (Figure 3d, panels for patient 25, and Extended data figure 2d). This same phenomenon was seen in patient 92 where pre-treatment cluster 1 had a similar expression pattern to a post-treatment cluster 2, suggesting there the IFN/IFM enriched pre-treatment tumor cells are in a similar phenotypic state as the post-treatment, “persister” cells. In addition, although both post-treatment clusters in sample 25 displayed elevated IFN and TNFA/NFKB signature scores relative to pretreatment clusters, they expressed different subsets of IFN and TNFA/NFKB genes. Similar differences were found in the other tumor samples, raising the possibility that there may be different transcriptional regulatory mechanisms driving these signatures in the different post-treatment populations.

The Proliferation-associated signatures were enriched in distinct clusters, supporting the anti-correlation with IFN/IFM signatures as shown in Figure 2. The EMT signature showed variable association with IFN/IFM signatures in most clusters, consistent with the variable correlation with IFNγ observed in Figure 2.

One question is whether the correlated programs are driven by specific genetic alterations. We considered it unlikely that correlated programs are driven by specific genetic alterations because the post-treatment enrichment of the IFN/IFM signatures was detected in all but one patient; however, we generated heatmaps of the inferred copy number alterations (CNAs) from the scRNA-seq data in the pre-post treatment tumor pairs (Extended data figure 3) to address this. Typical CNAs associated with ovarian cancer^4^ were observed across all patients, including gains of 3q or 8q, and deletions of 6q or 16p/q. There was enrichment or loss of specific CNAs post treatment unique to individual patients, for example 19q deletion (enriched in patient 25 post clusters). Some CNAs were heterogeneous across both pre and post clusters of the same patient (i.e. chr19 gain in patient 92 clusters 0, 1, 2 vs 3, 4). However, we did not find recurrent CNAs that could explain the observed enrichment of these signatures.

### IFN/IFM signatures are enriched in subpopulations of pre-treatment cancer cells across HGSOC tumors from large datasets

We examined three additional datasets to evaluate whether similar sets of correlated IFN/IFM programs are detectable in different cohorts of treatment-naive HGSOC; these included: 1) scRNAseq data that we generated from laparoscopic tumor biopsies (n = 37 patients) from recently diagnosed HGSOC patients enrolled in the Ovarian Cancer Precision Medicine Initiative (OCPMI) at the University of Minnesota (UMN OCPMI); 2) snRNAseq that we generated from frozen HGSOC tumor specimens (n = 53 patients) from the TRIO14^40^ clinical trial, and 3) HGSOC tumor specimens (n = 41 patients, n = 127 tumor samples) from the Memorial Sloan Kettering Cancer Center SPECTRUM cohort (MSK SPECTRUM^20^).

As observed in the matched pre-post tumor pairs, the correlated IFN/IFM a Proliferation-enriched gene signatures (Figure 4a,b) in the pre-treatment tumor cell clusters from the UMN OCPMI dataset similar to that described above in pre- vs post-treatment clusters (Figure 3c). In this cohort, IFM signatures were also correlated with IFNγ signatures in all samples, and anti-correlated with proliferation-related signatures (Figure 4c,d). In a second validation dataset using frozen samples from theTRIO14 study, we saw correlated signatures in the tumor cell clusters (Supplemental data Figure 2a,b) and correlated expression of IFM signatures with IFNγ and anti-correlated expression of proliferation-related signatures (Supplemental data Figure 2c,d). Although the Pearson correlations in the TRIO14 dataset were in the same direction, they were generally lower than seen in the UMN OCPMI dataset. The scRNA-seq data in the TRIO14 dataset was generated using frozen nuclei sequencing, while the UMN OCPMI dataset was generated using fresh whole cells (see Methods), which may account for this phenomenon.

**Figure 4:**
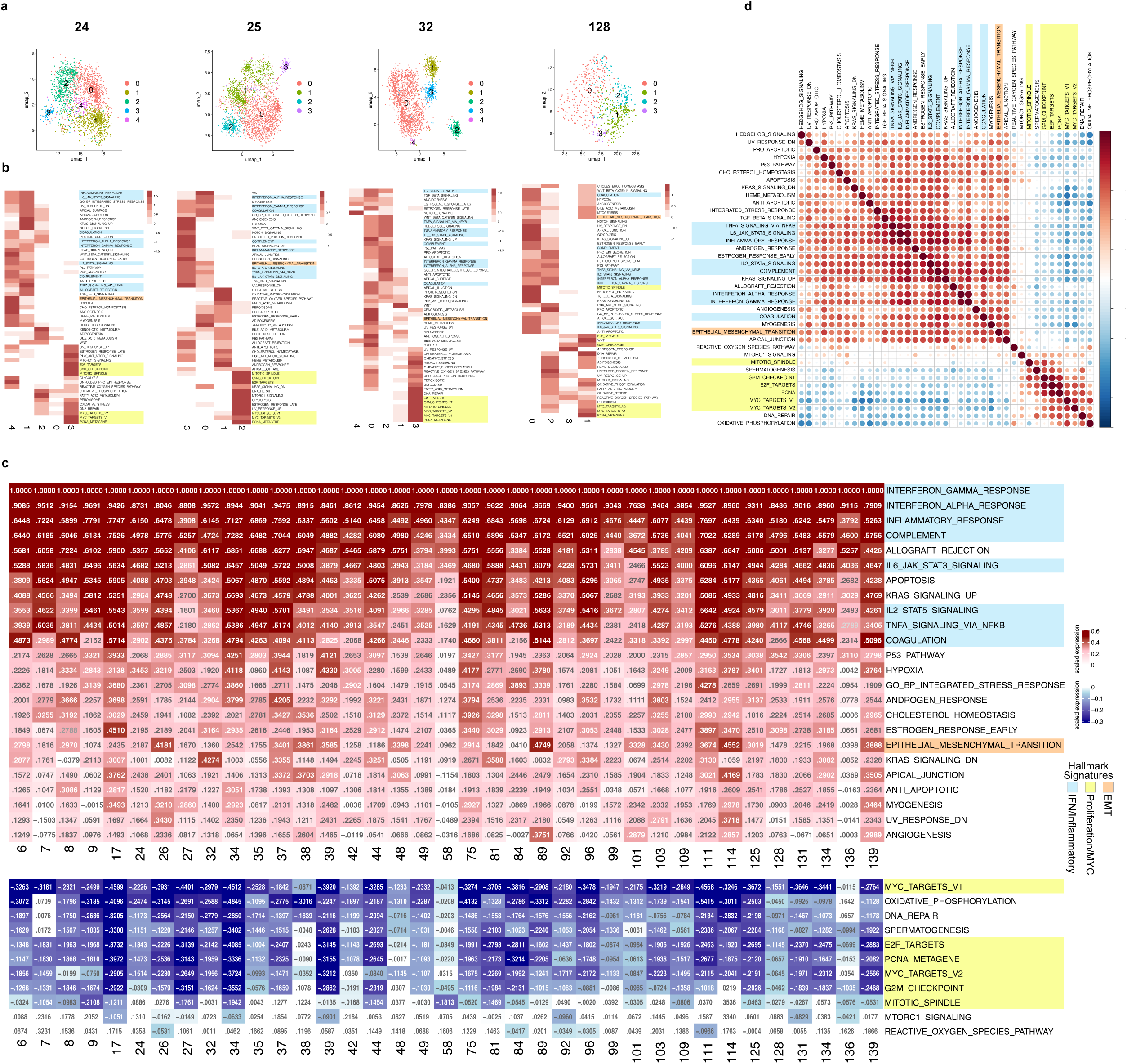
Analysis of pre-treatment tumors from UMN OCPMI dataset. a. UMAP plots of epithelial cancer cell clusters in four representative pre-treatment patient tumor samples color-coded by cluster. b. Heatmaps showing the signature scores of pseudo-bulk average expression of each cluster. c. Table showing the Pearson correlation for each signature compared to the IFNγ signature in each sample. The upper table shows the results for signatures that were found to be positively correlated with IFNγ and the lower table, those that were negatively correlated. Highlighting was used to highlight the different levels of correlation. d. Matrix showing the pairwise Pearson correlation values based on GSEA scores. Gene sets are colored by functional category as described in Figure 2.

In the UMN OCPMI and TRIO14 study, most of the samples were taken from the omentum. To evaluate whether a similar pattern of correlated signatures was observed in tumors from metastatic sites other than the omentum, we examined signatures correlated with IFNγ in the tumors from different sites associated with the MSKCC Spectrum patient collection (Extended data figure 4a-c). Across the whole dataset, tumors from different sites showed a similar set of correlated and anti-correlated signatures as observed in all of the other datasets.

Interestingly, the EMT signature was highly correlated (Pearson correlation > 0.4) with the IFNγ signature in less than 10% of samples in all three studies (3 of 37 UMN OCPMI, 4 of 53 TRIO14, 7 of 127 MSK SPECTRUM) (Figure 4c, Extended data figure 4c, Supplemental data Figure 2c). Furthermore, in the MSK SPECTRUM dataset there was notable variation within the different sites in the extent to which the EMT signature correlated with the IFN/IFM signatures suggesting that different sites might vary in the extent to which an EMT program is promoted (Extended data figure 4c).

To detect differences in expression patterns in different anatomical sites, we generated integrated datasets for patients with multiple tumor sites (Extended data figure 5). We detected two patterns of clusters in these patient tumors. In approximately half of the patients, tumor cells from each site were well-represented in each cluster from the integrated UMAP, indicating that tumor cells at different sites expressed similar transcriptional states (representative sample in Extended data figure 5a-c). In the other half, there was differential enrichment of cells in different clusters from each metastatic site (representative sample in Extended data figure 5d-f). To evaluate whether these distinct cluster patterns were associated with any differences in the correlated IFN/IFM signatures and proliferation/MYC signatures, we generated heatmaps from each of the different sites for each of the tumors in the second group (representative sample shown in Extended data figure 5g). While the tumors at different sites displayed distinct dominant clusters, each of the sites showed a similar pattern of correlated IFN/IFM- and proliferation-related signatures (Extended data figure 5h), suggesting that both cell states were represented at each site despite differences in the transcriptional clusters and tumor microenvironments.

To evaluate variation in the proportion of cells that are enriched for the IFN signature in tumors from different patients, we integrated the data from each patient across the three pre-treatment datasets and then calculated the IFNγ signature score, as a proxy for the IFN/IFM population, of each individual cell (Extended data figure 6a-c). The findings from all three datasets indicate that there is significant variation in the proportions of advanced metastatic HGSOC tumor cells that are enriched for IFN. Tumors with the lowest IFNγ signature scores were dominated by tumors that showed a Fold Back Inversion mutational signature, which is characterized by inverted duplications of DNA sequences and generally exclusive to cases that are homologous recombination defective (HRD)^50^.. The Shah laboratory has also found that tumors that contain prevalent cells with whole genome duplication express low levels of IFN-IFM signature genes^24^. These findings suggest that tumor mutational alterations can influence the activation of IFN and IFM programs: however, since half of the FBI tumors displayed much higher IFNγ signature score, other factors can contribute to the IFN-high phenotype

### Interferon and inflammation programs in the tumor microenvironment (TME)

Crosstalk between cancer cells and stromal cells in the TME influence tumor progression and evolution. We found a similar elevation of IFNγ and Inflammatory Response signature scores in post-treatment TME stromal cell types, including T cells, B cells, myeloid cells, fibroblasts, and endothelial cells (Figure 5, Supplemental data figure 3, Supplementary Table 6). Close examination of the individual IFNγ signature genes indicated canonical ISGs, indicative of IFN activation, were enriched in all types of stromal cells except in fibroblasts from tumor 25 and 92, where the canonical ISGs were not specifically enriched (Supplementary Table 6). A similar positive correlation between IFN module score and IFN/JAK-STAT pathway in T cells and macrophages was reported in the SPECTRUM data set^20^. Stromal cells from the patient with peritonitis (38) were highly enriched for the inflammatory response signature in the pre-treatment biopsy, as seen in tumor epithelial cells.

**Figure 5.**
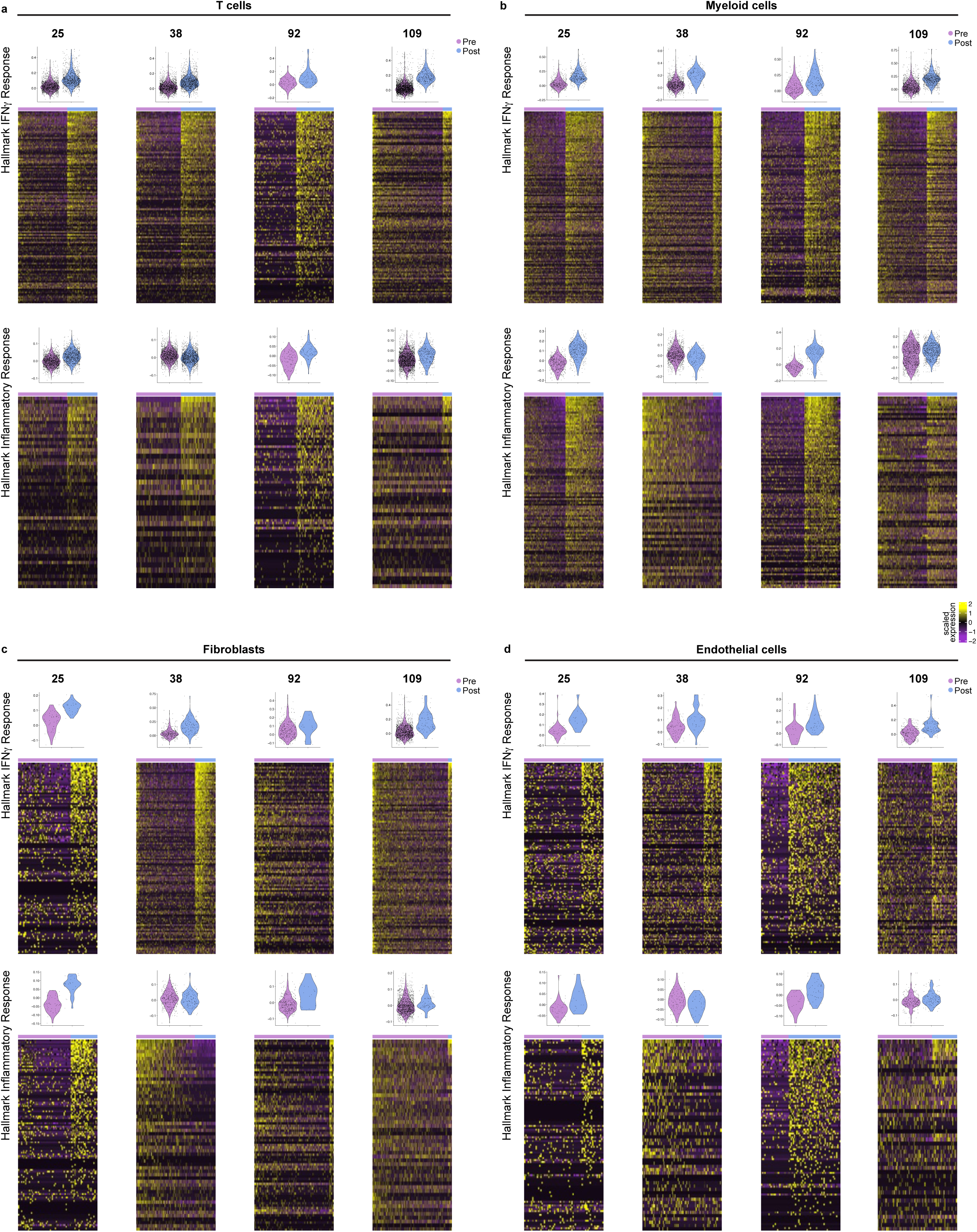
Analysis of Hallmark IFNγ and Inflammatory response signature genes in stromal cells from four patient tumor samples. Violin plots and gene expression heatmaps of the Hallmark IFNγ and Inflammatory response gene sets for patients 25, 38, 92 and 109 in (a) T cells, (b) myeloid cells, (c) fibroblasts, and (d) endothelial cells. Cells are grouped by pre-(purple) vs post-(blue) treatment in both the violin plots and the heatmaps and ordered by highest to lowest IFNγ score in each group in the heatmaps.

### Spatial analysis of matched tumor samples

The scRNAseq data above provide evidence that both epithelial tumor cells and stromal cells are enriched for IFN/IFM signature genes after treatment. IFN/IFM pathway activation in these cells could be due to paracrine signaling from one compartment to the other or driven by intrinsic cellular mechanisms. To address this question, we examined the spatial localization of ISG-positive (ISG^+^) cells and their local microenvironment utilizing multiplex cyclic immunofluorescence (CyCIF) imaging. This approach enabled us to evaluate whether IFN-induced genes were stably expressed at the protein level, and to evaluate IFN pathway activation in the two samples (patients 96 and 139) that had low cancer cell numbers in the scRNAseq analysis. Of particular interest was whether direct interactions between ISG^+^ tumor or ISG^+^ stromal cells contribute to the ISG-positive state of either population.

To analyze the spatial architecture of pre- and post-treatment tumors, we developed a CyCIF antibody panel and computational pipeline that classified over eleven million cells using cell-type markers for tumor cells, B cells, cytotoxic T cells, myeloid populations, endothelial cells, fibroblasts, and adipocytes (Supplementary table 7; see Methods). Additionally, CD45^+^ cells lacking expression of CD8, CD20, CD68, CD163, or CD11b were classified as immune cells not otherwise specified (Other Immune). There was considerable heterogeneity in tissue organization and cell type architecture across the samples (Figure 6a,b and Extended data figure 7a); in most samples, tumor cells, fibroblasts and myeloid cell populations were the dominant populations (Extended data figure 7b). In three samples, CD36^+^ adipose tissue from the underlying omentum was prominent (Pre-38,109, Post-96). We did not observe any consistent differences in cell type proportions between pre- and post-NACT tissues.

**Figure 6.**
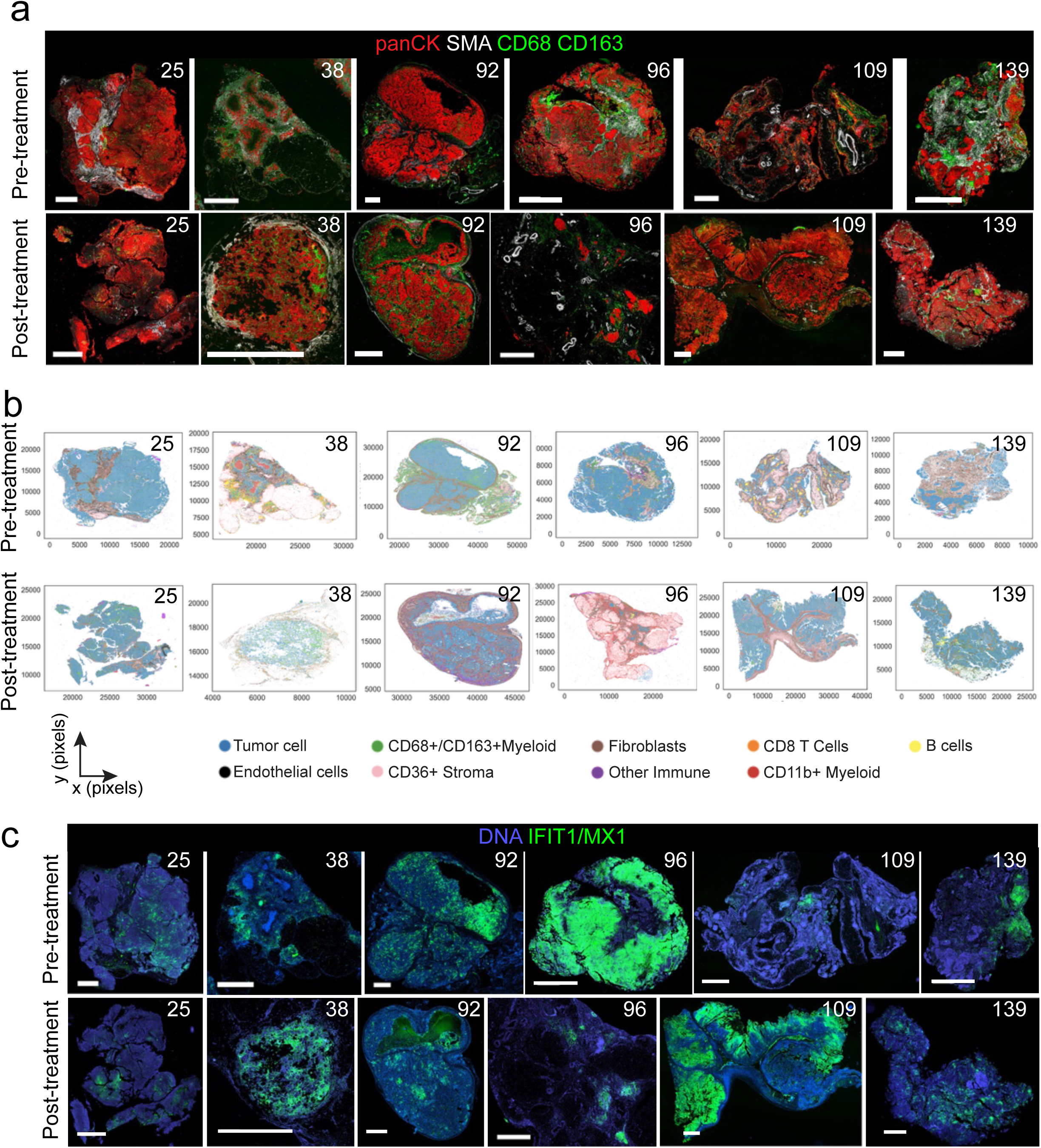
HGSOC samples exhibit diverse architecture, but spatially clustered IFIT1 and MX1 expression. a. CyCIF images showing pan-cytokeratin (red), smooth muscle actin (SMA, white), and CD68 and CD163 (green) in patient tumor pre-treatment (top row) and post-treatment (bottom row) samples. b. Centroid plots of tumors from (a) showing the results of cell type classification. c. CyCIF images showing IFIT1 and MX1 (green) staining in same tumors as in (a). All scale bars are 2 mm.

To identify IFN pathway-active cells, we quantified the combined staining intensity scores for IFIT1 and MX1 which were commonly expressed in tumor cells and were positively correlated with the IFNγ response signature score based on scRNAseq data (see Methods).

IFIT1 and MX1 staining was discrete and focal in the majority of samples (Figure 6c, Extended data figure 7c) except for the pre-treatment sample from patient 96 and the post treatment sample from patient 109, in which the ISG^+^ areas were broadly distributed across the entire tissue. The abundant expression of these proteins in the pre-treatment tissue from patient 96 agreed with the observations from our scRNASeq analysis in which there was a markedly high proportion of ISG^+^ tumor cells pre-treatment (Supplemental data figure 4,). The post-treatment tissue from patient 96 contained multiple small, spatially-separated tumor nodules, most of which were ISG^+^, thus mimicking the pre-treatment tissue with abundant ISG^+^ tumor cells. To quantify the clustering of ISG^+^ cells, we applied the spatial interaction function in SciMap^41^ to measure ISG^+^ cell interactions with each other or ISG-negative (ISG^-^) cells. Across all of the tissues, cells of the same ISG status associate strongly with themselves irrespective of cell type identity (Extended data figure 7d), creating zones of ISG^+^ and ISG^-^ cells with very little mixing of these two states. A similar level of ISG^+^ cell clustering was also observed when this analysis was performed on tumor cells alone, indicating their strong spatial association with each other (Extended data figure 7e).

To define the cell populations that express ISGs, we quantified the ISG^+^ cells in each sample. In most patient samples (11 of 14), tumor cells represented the largest fraction of ISG^+^ cells (approximately 30-80%; Figure 7a). Myeloid cells, fibroblasts and endothelial cells generally ranged from 8-47% of ISG^+^ cells. We also detected ISG expression in T cells and other stromal cells, but these represented small fractions of the total ISG^+^ cell population.

**Figure 7.**
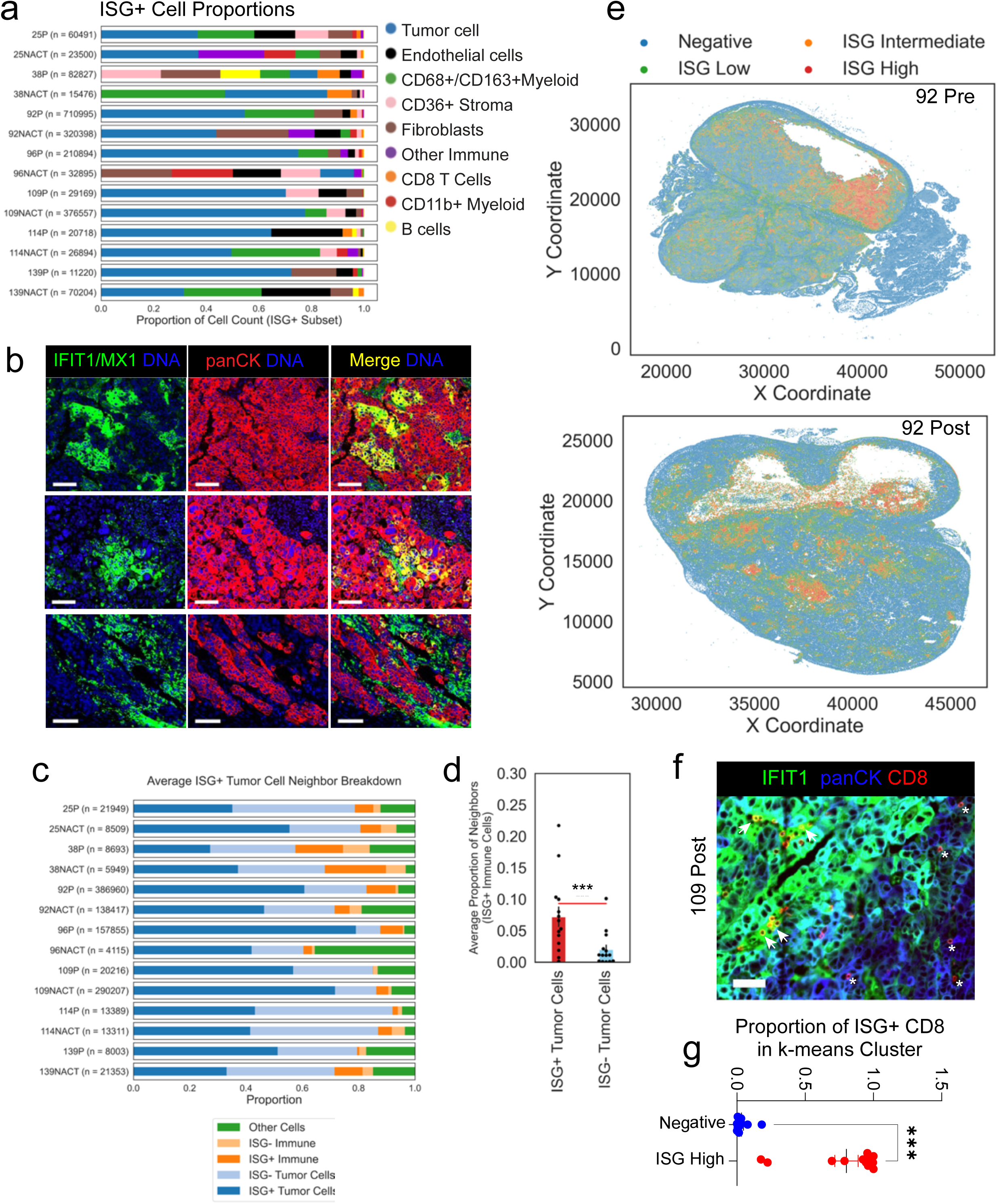
Spatial analysis of ISG-positive cells. a. Stacked bar plot showing proportions of ISG^+^ cell type across tumors. The total number of cells per sample is shown in parentheses. b. CyCIF images showing the expression of IFIT1 and MX1 (green) in tumor (panCK^+^, red) and neighboring stromal cells. DNA is pseudo colored in blue and tumor cells expressing ISGs appear yellow in the merged images, while non-tumor cells expressing ISGs are green. Each row is representative of the three patterns observed, using patient 92 tissue as an example (see text). All scale bars are 100 µm. c. Stacked bar plot showing the proportions of ISG^+^ tumor cell neighbors. Blue are tumor, orange are immune, and green are other cells. Darker shade indicates ISG^+^ cells (see legend). Total queried ISG^+^ tumor cells are shown in parentheses. d. The average proportion of neighbor ISG^+^ immune cells to either ISG^+^ tumor cells (red bar) or ISG^-^ tumor cells (blue bar) were compared using Wilcoxon’s signed-rank test. e. Visual representation of the four k-means clusters of all cells for IFIT1 and MX1 expression (negative=blue, ISG^low^=green, ISG^Interm^=orange, ISG^high^=red) were mapped spatially as centroid plots. f. CyCIF image showing CD8 (red) T cells in and around an IFIT1^+^ (green) cell cluster. Tumor cells (panCK^+^) are pseudocolored blue. Scale bar = 50 µm. Arrows highlight IFIT1^+^ CD8 T cells, asterisks show IFIT1^-^ CD8 T cells. g. The proportion of CD8 T cells in the Negative or ISG^high^ clusters that express ISGs were compared using Mann-Whitney. Throughout figure, *** indicates p<0.001.

Having established the diversity of ISG^+^ cell types and their propensity to form cohesive zones, we viewed the ISG^+^ areas at higher magnifications to assess their cell type composition. Overall, there were three distinct “cell neighborhoods”: (1) tumor-rich, stroma-free areas harboring ISG^+^ tumor cells clustered together (Figure 7b, top row), (2) ISG^+^ stromal cells in close proximity to ISG^+^ tumor cells, (Figure 7b, middle row), and (3) ISG^-^ tumor cells adjacent to ISG^+^ stromal cells (Figure 7b, bottom row). Notably, ISG^+^ tumor cells were not restricted to areas with extensive ISG^+^ stromal cell infiltration and were frequently noted in tumor-dense, stroma-free areas.

To quantitatively describe the cells neighboring ISG^+^ tumor cells, we counted the different cell types within a 20-micron radius of each ISG^+^ tumor cell. In all tissues, the majority of ISG^+^ tumor cell neighbors were other tumor cells, either those expressing or not expressing ISGs) (Figure 7c, Extended data figure 7f). In most tissues, the most common stromal cell neighbors of ISG^+^ tumor cells were endothelial cells and CD68^+^/CD163^+^ myeloid cells (Extended data figure 7f). CD8^+^ T cells expressing ISGs represented a small proportion of ISG^+^ tumor cell neighbors. These analyses support the conclusion that ISG^+^ tumor cells are predominantly found adjacent to epithelial tumor cells, providing evidence that activation of the interferon pathway in most tumor cells is mediated by tumor intrinsic mechanisms. Although immune cells were not a major neighbor of ISG^+^ tumor cells, the ISG^+^ subset of immune cells preferentially associated with ISG^+^ tumor cells (Figure 7d).

While the neighbor analyses provided insight into directly adjacent cells of ISG^+^ tumor, it did not capture larger ISG^+^ zones like those visible in the immunofluorescence images in Figure 6c. Thus, we sought to computationally define the ISG^+^ zones using only MX1 and IFIT1 and evaluate the cell type composition of these spatial neighborhoods by performing k-means clustering on all cells in the tissues – except for 109P and 139P which were excluded due to small numbers of ISG^+^ T cells – and visualizing the four resulting clusters in the tissue (Figure 7e, Extended data figure 8a-b). In general, clustering separated cells based on MX1 and IFIT1 intensity, with the highest staining of both detectable in a cluster termed ‘ISG^high^’ and cells with the lowest levels of both markers in ‘ISG^low^’. We found that the ISG k-means clusters were distributed in a distinct spatial pattern. Specifically, the ISG^high^ and ISG-Intermediate (ISG^Interm^) clusters formed foci that were surrounded by the ISG^low^ cluster (Figure 7e, Extended data figure 8b) suggestive of an IFN gradient established by its secretion and subsequent dispersion as has been previously noted^42^. This result excludes the possibility that these cells are randomly mixed, instead showing a spatial gradient of ISG expression.

The ISG^high^ clusters were predominantly composed of tumor cells, with CD68^+^/CD163^+^ myeloid cells being the next most well-represented cell type (Extended data figure 8c). Fibroblasts and endothelial cells showed variable representation from 0-40%. In all but two tissues, CD8^+^ T cells represented fewer than 10% of cells in the ISG^high^ zones and their proportions were not significantly different between ISG^high^ and Negative zones (Extended data figure 8c). While T cells represent a small population of cells in the ISG^high^ clusters, IFNγ is known to enhance the cytotoxicity of T cells and thus analysis of ISG^high^ T cells is important in assessing tumor immune status. In this dataset, ISG^+^ T cells co-express markers of cytotoxic T cells (Extended data figure 8d-g), and T cells are the predominant IFNγ expressors (Extended figure 8h-l).Visual inspection of the ISG^high^ zones provided evidence that the CD8^+^ T cells in these areas were predominantly ISG^+^, whereas those outside the ISG^high^ zones were ISG^-^ (Figure 7f). Quantification of the ISG^+^ T cell proportions in the ISG^+^ and negative zones strongly supported this observation; almost all of the CD8^+^ T cells found in the ISG^high^ k-means clusters were ISG^+^ (Figure 7g). The comparable cell type composition between these two zones (Extended data figure 8c) agreed with earlier results supporting the conclusion that ISG expression is not restricted to areas of immune infiltration or tumor-stroma interaction.

Our analyses demonstrates that ISG^+^ cells spatially associate with each other more than with ISG^-^ cells (Extended data figure 7d). Areas of ISG^+^ cells were found to be composed entirely of tumor cells or have a mix of tumor cells and diverse stromal cell types (Figure 7b, f), indicating that there is no specific cellular environment that is necessarily associated with ISG expression. However, there is evidence of ISG^+^ CD8 T cell association with ISG^+^ tumor cells and of enrichment of ISG^+^ CD8 T cells in ISG^high^ zones within these tumors (Figure 7g). Collectively, these results suggest that tumor cell-intrinsic mechanisms play an important role in ISG expression, though paracrine mechanisms may also contribute, and that CD8^+^ T cells expressing an activated interferon pathway are highly enriched in the ISG^+^ tumor zones.

### Spatial analysis of HGSOC precursor lesions

Having identified this tumor cell population with a correlated IFN/IFM program and found that IFN pathway activation is predominantly tumor cell-intrinsic, we addressed whether this population is present in the precursor lesions of HGSOC. A recent report showed that IFNα and IFNγ mRNA signatures are upregulated in STIC lesions relative to p53-signature fallopian tube epithelial cells^43^. Furthermore, the authors observed a trend of increasing expression of this signature over the course of tumorigenesis, suggesting an early induction of the IFN/IFM state and prompting us to examine the spatial localization and ISG expression in STIC lesions by staining a set of 11 tissues from patients with HGSOC that had confirmed STIC lesions with a modified CyCIF panel (Supplementary Table 7). We found expression of the ISG marker IFIT1 in patches of cells across all STIC lesions (Figure 8a). Furthermore, these ISG^+^ epithelial cells were often not adjacent to ISG^+^ stromal cells, (Figure 8a; asterisks show individual immune cells, with yellow being ISG^+^, arrows point to IFIT1^+^ epithelium without neighboring immune cells) indicating that the epithelial cells are not reliant on stromal cells for IFN pathway activation and ISG expression.

**Figure 8.**
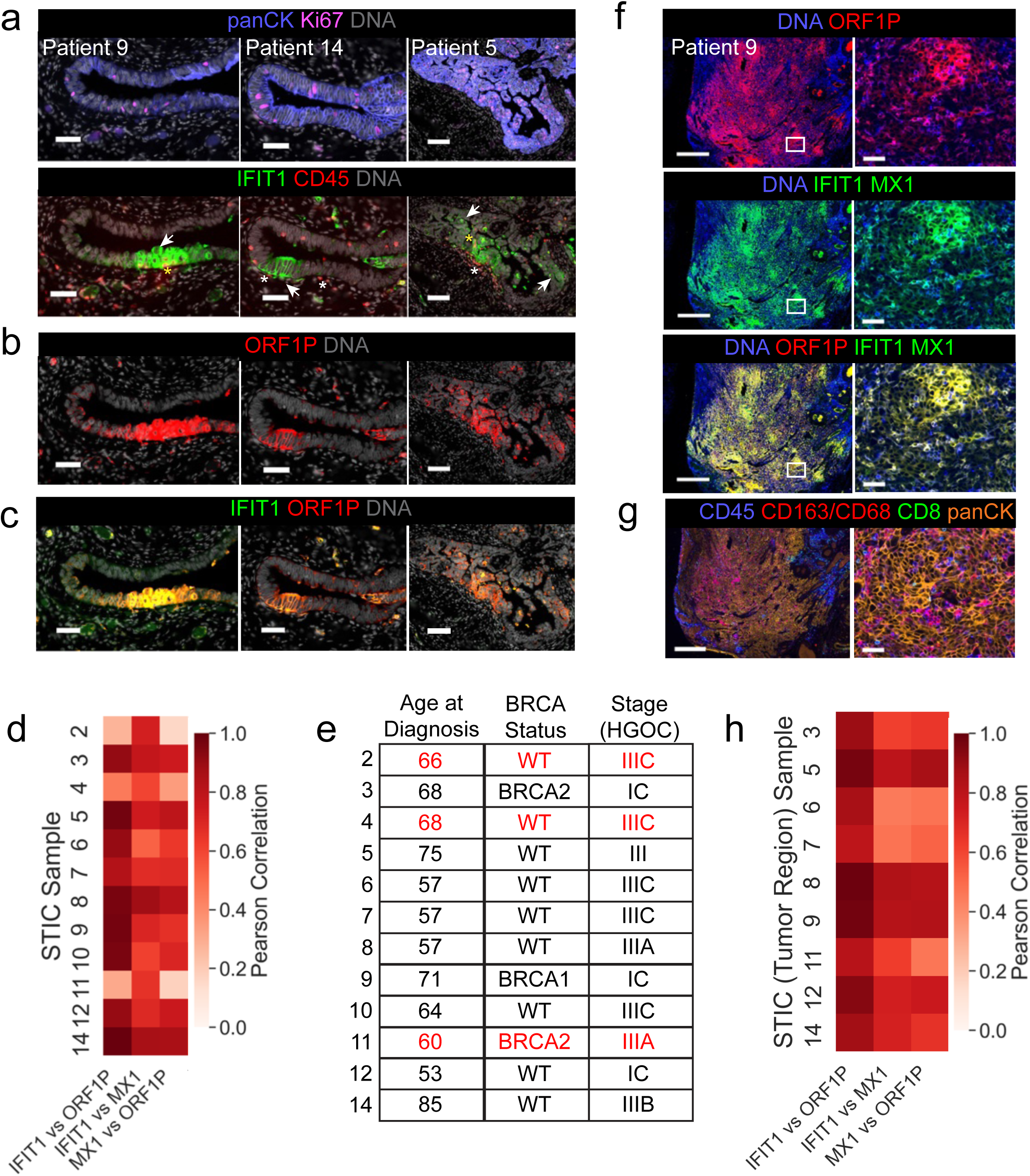
Spatial analysis of ISG-positive cells in STIC lesions and associated HGSOC. a. CyCIF staining of highly proliferative (Ki67^+^, magenta), panCK^+^ (blue) STIC lesions. IFIT1 expression (green) and immune cell (CD45^+^, red) presence in these areas are shown in the bottom row. Asterisks denote individual immune cells (yellow = ISG^+^), arrows point to IFIT1^+^ epithelium without neighboring immune cells. b. CyCIF staining of ORF1P (red) in the same STIC lesions as shown in (a). c. CyCIF staining showing colocalization (yellow) of IFIT1 (green) and ORF1P (red) in STIC lesions. All scale bars in a-c are 50 µm for patients 9 and 14, and 100 µm for patient 5. DNA is colored gray. d. Heatmap showing the single-cell level correlation of indicated markers in STIC epithelial cells. e. Patient age at diagnosis, BRCA mutation status, and HGSOC grade. Red font highlights patients in which ORF1P and ISGs showed a weaker correlation. f. CyCIF staining of HGSOC present in tissues with STIC lesions showing ORF1P (red), IFIT1 and MX1 (green), DNA (blue), and the overlap of IFIT1 with ORF1P (yellow). Boxed area in left hand panels (scale bar = 50 µm) is magnified on right (scale bar = 500 µm) g. CyCIF staining of the same areas as in (f) showing tumor (panCK, orange), CD8 T (green), myeloid (CD68 & CD163, red) and immune cell (CD45, blue) distribution. Scale bars as in (f). h. Heatmap showing single-cell level correlation of indicated markers in tumor epithelial cells.

IFN programs can be induced by de-repression of repetitive elements (REs) in the genome, and REs are commonly chronically expressed in many types of tumors^44–46^. STIC lesions have been shown to express ORF1P, a protein product of the repeat element, LINE1^47,48^. Thus, the presence of LINE1 and other REs in the STIC epithelium may account for the apparent cell-intrinsic activation of the IFN pathway in these lesions. Visual inspection confirmed that ORF1P was expressed in STIC lesions, but not normal epithelial cells (Figure 8b & Extended Figure 9a) and predominantly co-localized with IFIT1 (Figure 8c). To quantify this relationship, we computationally extracted STIC lesions from our CyCIF images for analysis and performed single cell segmentation yielding over 80,000 epithelial cells (a median of approximately 5000 epithelial cells per patient). A correlation analysis of the markers IFIT1, MX1, and ORF1P in pan-keratin-positive cells shows that in eight of the eleven patients, these markers correlated strongly with each other, with IFIT1 correlating especially strongly with ORF1P (Figure 8d). Neither age of diagnosis, BRCA status, nor cancer stage explain the low correlation in the three outlier patients (Figure 8e, red font highlights poorly correlated patients).

The observation that ORF1P and ISGs are co-expressed in STIC lesions raised the question whether these two proteins are also co-expressed in tumor cells, and as such, would provide a potential mechanism for cell-intrinsic IFN pathway activation. Nine of the STIC tissue sections had tumors present. Visually, these tumors showed remarkable overlap of IFIT1 and ORF1P (Figure 8f) and were not necessarily located in stroma-rich areas (Figure 8g, Extended figure 9b). Computational extraction of these tumor regions and quantification of this relationship at the single cell level confirmed strong correlations between IFIT1 or MX1 and ORF1P (Figure 8h). Thus, our spatial analysis of these tissues provides evidence that derepression of repeat elements like LINE1 is a likely cause of IFN pathway activation and subsequent ISG expression, and that the IFN/IFM state is induced in the early stages of disease.

## Discussion

Single cell RNA analyses of more than 130 HGSOC samples identified a subpopulation of tumor cells that are highly enriched for interferon pathway genes and a set of correlated inflammatory programs. Expression of these programs was anti-correlated with proliferation-associated programs and MYC-targets, while tumor cells enriched for MYC-targets and proliferation-related programs were associated with cell clusters transcriptionally distinct from those enriched for IFN/IFM signatures. The distinct IFN/IFM- and Proliferation/MYC-enriched populations were detected using either scRNAseq or snRNAseq platforms in three different patient cohorts and in different metastatic sites, indicating that the existence of these two populations is a feature of HGSOC. Previous studies involving NMF and DGE analyses of cellular programs enriched in treatment-naive HGSOC have also identified cell populations enriched for interferon and/or JAK/STAT pathway programs^16,20–23^; however, in only one study did they note the set of correlated programs. Nath and coworkers used a computational approach to define three overarching archetypes of HGSOC, two of which (‘Metabolism and Proliferation’ and ‘Cellular Defense Response’) are highly similar to the two major populations defined in our study, and highly anti-correlated with each, as in our study^36^. The in-depth analysis of the IFN/IFM programs in individual tumor samples in our study provides a deeper characterization of genes associated with the IFN/IFM programs in tumor and stromal cells that is not feasible with integrated datasets from many tumors.

An important finding from this study is that IFN/IFM programs were strongly enriched in tumor cells which survived chemotherapy in six of seven tumors. Thus, this enrichment is likely not associated with a specific tumor genotype, but is more likely due to selection of cells in drug tolerant states and/or adaptation to chemotherapy. The evidence for a similar enrichment of IFN/IFM response programs in stromal cells suggests that chemotherapy induces an inflammatory state throughout the diverse ecosystem of HGSOC. However, the evidence that the proportion of IFN-high cells in pre-treatment samples correlated with specific mutational signatures suggests that the baseline IFN/IFM state is influenced by genotype (Extended data figure 6). Only a few studies of scRNAseq analysis of matched pre- and post-treatment HGSOC samples have been reported. Comparison with our study is difficult because of the highly variable set of samples, distinct focuses of the studies, and the common approach to integrate all samples for initial transcriptional cluster analysis. We opted to analyze sample by patient which avoided clustering based on potentially misleading clustering due to stronger transcriptional similarities with subset of genes in other tumor samples. One study reported post-treatment enrichment of a ‘stress program’^33^ that shares many genes with the Hallmark TNFA/NFKB signature, especially IEGs which dominated the DEGs in this signature in our study (Supplementary Table 3). Pseudo-bulk analysis of integrated patient samples in another study showed TNFα and EMT signature enrichment post-treatment^34^. We detected enrichment of EMT in half of our matched pairs, and cells with the EMT signature correlated with the IFN-IFM signatures in a subset of tumors across all three cohorts. Chemotherapy has been shown to enrich for IFN-IFM populations in other types of cancer^49–52^.

Spatial analysis using multiplex immunofluorescent imaging indicated that interferon pathway-activated tumor cells (ISG^+^ cells) were generally localized in clusters. While the majority of ISG^+^ tumor cells were not directly surrounded by ISG^+^ immune cells and thus, not likely to be activated through paracrine interactions, ISG^+^ tumor cells were typically the predominant cell type in larger ISG^+^ zones that contained multiple ISG^+^ stromal cell types. In contrast, ISG^+^ immune cells were more likely neighbors of ISG^+^ tumor cells than ISG^-^ tumor cells, raising the possibility of paracrine activation of immune cells by IFN produced by ISG^+^ tumor cells. Since the multi-cellular ISG^+^ zones did not display evidence of necrosis, they are likely not areas rich in dying cells which release Damage-Associated Molecular Patterns (DAMPs) that would recruit and activate immune cells. A previous CyCIF cell proximity analysis of HGSOC tumor sections showed that tumor cells expressing MHCII proteins, which are known ISGs and commonly correlated with ISGs in our and others’ analyses, frequently associate with immune cells and are located in tumor-stroma interface regions where the authors postulated they coordinated an immune response^25,53^. Similarly, spatial transcriptomic analysis of HGSOC tumors have shown that tumor cell neighbors of ISG^+^ tumor infiltrating lymphocytes (TILs) are highly enriched for ISGs, supporting the spatial association of ISG^+^ tumor cells and IFN-active TILs.

An important question raised by these studies is: what drives the induction of the interferon pathway? The spatial analysis provided evidence that adjacency of tumor cells with ISG^+^ stromal cells cannot explain the majority of ISG activation in tumor cells; however, the highly localized adjacency of some ISG^+^ tumor cells to ISG^+^ stromal cells suggest that paracrine IFN induction occurs in some tumor regions. Activation of ISGs by cytoplasmic DNA associated with DNA damage was considered as another possible mechanism to explain the chemotherapy-induced ISG enrichment; however, we did not detect induction of any DNA damage repair signatures, nor a correlation of ISGs with phosphorylated H2AX or micronuclei in pre- or post-treatment tumors. While DNA damage responses are commonly activated by chemotherapy, this acute response was likely not sustained in the tumors during the minimal four-week interval between chemotherapy treatment and interval debulking surgery. A third possible mechanism for ISG induction in tumor cells is through dsRNAs or dsDNA generated from depression of endogenous retrotransposons and repeat RNAs^44–46^. This possibility is strongly supported by our evidence that ISG expression in early HGSOC precursor lesions, STIC lesions, and fallopian tube-associated primary invasive HGSOC is highly correlated with the expression of ORF1P, a protein product of LINE1 retrotransposons. Reports from others have shown that LINE1 DNA is demethylated in STIC lesions^54^, that deletion of p53, which is known to suppress derepression of repetitive elements^55–57^, can induce REs in FTEs^58^, that ORF1P is expressed in STIC lesions^47,48^, and that interferon signatures are expressed in STIC lesions^43^; however, this study provides the first evidence that the ISG expression is highly correlated with LINE1 ORF1P protein expression, thus linking derepression of this RE with interferon program activation.

The anti-correlation of MYC-target signatures with triple negative breast cancers IFN/IFM gene signatures in all three HGSOC cohorts is consistent with multiple reports showing MYC suppression of REs and ISGs in multiple models including MYC binding to promoters in young LINE1 retrotransposons and anti-correlation of LINE1 and MYC expression in TCGA ovarian tumors^59^, MYC suppression of IFN in tumors induced by deletion of Brca1 and Tp53^60^ and in TNBCs^61–63^.

A model consistent with our findings and published reports^25,58^ is that early events in transformation of FTEs promote demethylation of REs through previously reported mechanisms, i.e. loss of Rb^64,65^, alterations in methyltransferases or demethylases^54^ or H3.3 chaperone complex genes^66^). Prior mutation of p53, the presumed first event initiating transformation of FTEs, allows expression of the RE RNAs^55–57^. RE RNAs can suppress tumor progression through induction of interferon which have been shown to suppress proliferation and induce senescence, intrinsic programmed cell death and secretion of pro-apoptotic paracrine signals in tumor epithelial cells, and also promote immune cell activation and clearance of initiated cells^67–69^. Escape from RE RNA-mediated tumor suppression could be mediated by a variety of different mechanisms, including but not limited to, high MYC expression, ADAR modification of RE RNAs^70^, high ORF1P expression which can act as a sponge of ALU inverted repeats RNAs^70^, and RAS/AKT signaling which suppresses DDX60 and other interferon-stimulated genes^71^. Many questions remain to be answered relating to this model in order to understand (1) what triggers RE derepression, (2) whether STIC-associated ISG/ORF1P-negative cells have escaped RE-mediated suppression, (3) how the quiescent ISG^+^ cells within invasive tumors are maintained, (4) how dynamic is the ISG^+^ state, (5) what mechanism is responsible for their enrichment after chemotherapy, and (6) how do these cells influence immune clearance of tumor cells.

While our study provides evidence that tumor cell populations that survive chemotherapy are highly enriched for IFN and IFM programs, the specific functional effects of interferon on cancer therapy resistance is an open question. In tumor models, interferon has been shown to either directly inhibit tumor cell growth and enhance tumor immune response, or to contribute to tumor therapy resistance and immune escape^49,72–77^. This variation in functional effects is in part related to acute^78,79^ versus chronic^80–82^ effects of interferons, but also due to both pro- and anti-inflammatory effects of ISGs. Elucidation of the functional consequences of this program on chemotherapy responsiveness will require further studies in tractable model systems. Regardless, our study suggests that targeting the proportionally large IFN-IFM cell populations that survive chemotherapy should be explored as a strategy to delay or prevent relapse after chemotherapy.

## Methods

### Patient enrollment and sample acquisition

All patient samples from the OCPMI UMN cohort were collected using protocols approved by the University of Minnesota’s Institutional Review Board (Protocol numbers 1408M52905 and 1611M99903). Women were recruited if they were at least 18 years of age and had suspected ovarian, primary peritoneal, or fallopian tube cancer based on imaging and biomarker tests. Final consent and enrollment occurred after confirmation of a histologic diagnosis of ovarian cancer based on a surgical sample. Multiple samples from the primary tumor specimen were collected in media on ice at the time of surgery and samples were immediately transported to the laboratory for processing. Fresh tumor tissue was divided into sections and were processed by: 1) flash freezing for DNA, RNA, and protein extraction, 2) fixed in formalin for histology, and 3) minced and placed in dissociation media for single cell sequencing.

Frozen patient samples and clinical data from the TRIO14 clinical trial were collected by the clinical trial study team^40^ and were shipped to the University of Minnesota for processing and sequencing.

### Clinical data

Clinical data, including pathologic diagnosis, stage, grade, MRD, BRCA status, PFS, OS, and chemotherapy response was extracted from the patient’s EMR for the UMN OCPMI cohort. Chemotherapy Response Scores (CRS) were assigned by a Molecular Pathologist (AN) based on published guidelines^83^.

### scRNAseq sequencing

Fresh biopsies collected from the UMN OCPMI cohort were sequenced at the University of Minnesota’s Genomics Center using the 10x Genomics Single Cell 3’ Protocol utilizing the ChromiumTM Single Cell 3’ Library & Gel Bead Kit and ChromiumTM Single Cell A Chip Kit. Approximately 5000 cells were partitioned into nanoliter-scale Gel Bead-In-EMulsions (GEMs) with one cell per GEM. Within each GEM, cells were lysed, primers were released and mixed with cell lysate. Incubation of the GEMs produced barcoded, full-length cDNA from mRNA. The full-length, barcoded cDNA was then amplified by PCR prior to library construction. Sequencing was performed using an Illumina HiSeq 2500 or NovaSeq.

Frozen samples (50-100 mg each) from the TRIO14 clinical trial were homogenized using a glass dounce tissue grinder containing 250 µl of ice cold EZ prep lysis buffer (Sigma-Aldrich). Sample was transferred to a 1.5 ml microcentrifuge tube, incubated on ice for 5 minutes and centrifuged at 500g for 6 minutes at 4°C. Pellet was resuspended in 1 ml lysis buffer and centrifuged a second time. Pellet was washed in 1 ml nuclei suspension buffer (Bovine serum albumin 0.02%, RNase Inhibitor 0.2U/ul, in 1xPBS), filtered through a 10µm cell strainer and counted. Single cell barcoded cDNA was generated using the 10X Genomics Chromium 3’ Library & Gel Bead Kit as described above. Sequencing was performed using an Illumina NovaSeq.

### scRNAseq analysis

#### Sequence mapping and quantification

Illumina raw sequencing output files were processed using Cell Ranger™ software using the human genome reference (GRCh38) to produce a filtered gene x cell matrix of UMI counts. Quality control measures included assessing gene and UMI counts, mitochondrial content, and other cell-level metrics to evaluate data integrity. The filtered matrix was used as input for downstream analysis using Seurat (v5.0.3)^84^ and SingleR (v2.4.1)^85^ using customized R (v4.3.2) scripts (available upon request).

#### Differential gene expression analysis

Differentially expressed genes were computed using edgeR (version 3.36.0). The pseudo-bulk gene expression was estimated by the average expression of each gene in Pre and Post groups. Thereafter, edgeR identified and ranked the differentially expressed genes between Pre and Post cell.

#### Cell clustering and annotation

Significant principal dimensions were determined by an elbow plot analysis. A resolution grid search ranging from 0.005 to 0.3 was conducted to identify cell type clusters, with annotation performed based on cell type-specific marker genes. Cancer cells were distinguished, and optimal cancer subclones were curated by finding cell cluster separation in the UMAP space per sample, employing a grid search on resolution (ranging from 0.05 to 2.00 with a step size of 0.05) and 150 replication parameters. Cell clusters were annotated with cell types by the cell type unique cell markers.

#### Copy number variation estimation

Inferred cancer subclonal copy number variation (CNV) scores were obtained using InferCNV (version 1.18.1), utilizing a minimum UMI count per gene (>0.1) and normal cells (other cell types) as the reference. Additional parameters considered during Infercnv analysis included the detection sensitivity threshold and statistical significance cutoffs for identifying CNV events.

#### Hallmark gene sets and hallmark enrichment scores

The 50 hallmark gene pathway sets were collected from MsigDB (version 7.2.1)^86^, and 8 additional gene pathways, i.e., ANTI_APOPTOTIC, PRO_APOPTOTIC, PCNA, GO_BP_INTEGRATED_STRESS_RESPONSE, ATF, OXIDATIVE_STRESS were self-curated from literature. The criteria for selecting hallmark gene sets included relevance to the biological processes of cancer study interest and robustness of gene annotations. The hallmark enrichment scores for individual cancer cells were estimated by the Seurat package AddModuleScore function with collected hallmark gene sets. The pseudo-bulk hallmark enrichment scores for cancer cell cluster was computed by using Escape (version 1.12.0)^87^, taking into account the pseudo-bulk average gene expression of each cell cluster by Seurat’s AverageExpression function.

Hallmark score and gene correlation within individual samples:

Given n cells, the scaled expression of gene X as (x_1_, x_2_, … x_i_, …, x_n_), and the hallmark of H scores as (h_1_, h_2_,, … hi, …, h_n_), the individual cell level Pearson correlation r between hallmark score H and gene X is as follows: 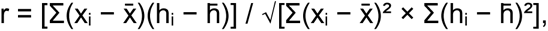 where 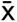 and 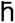 are the mean values of X and H respectively. Similarly, the individual cell level Pearson correlation r between hallmark H as (h_1_, h_2_,, … h_i_, …, h_n_) and hallmark K as (k_1_, k_2_,, … ki, …, k_n_) is as follows: 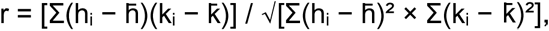 where 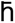 and 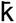 are the mean values of h and k respectively.

The Pearson correlation in multiple samples as the data set:

The Pearson correlation across samples considers the overall correlation among all pairs of samples clusters. Given one sample s_i_ with m_i_ clusters, the differential expressed genes of a pair of clusters were computed by Seurat’s FindMarkers function. The hallmark score of each pair of clusters were computed by fgsea’s (version 1.20.0)^88^. fgseaMultilevel function by the differentially expressed genes between this pair of clusters. The differentially expressed genes were collected by edgeR (version 3.36.0). The hallmark score of the sample S_i_ is h_si_ = (h_0_, h_1_, … h_i,_, h_l_), where l = mi⋅(mi−1)/2. By this means, the hallmark h’s score of all samples and hallmark k’s score of all samples are derived by the concatenation of all samples as h = (h_1_, h_2_,…, h_i_,… h_n_) and k = (k_1_, k_2_, …, k_i_, … k_n_) respectively. Thereafter, the pearson correlation r is defined as follows: 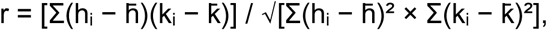 where 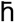 and 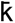 are the mean values of h and k respectively.

Jaccard distance between two hallmark gene sets:

Given two sets of hallmark genes G as (g_1_, g_2_, … g_i_, …, g_m_) and K as (k_1_, k_2_, … k_i_, …, k_n_) with m genes and n genes respectively, the Jaccard distance j is as follows: j = 1 − c / (m + n − c), where c = |G ∩ K|.

### Spatial Analyses

#### Tissue staining

For CyCIF, two serial, five micron FFPE tissue sections were manually dewaxed and rehydrated as previously described^89^. Next, the tissue sections underwent antigen retrieval using Antigen Unmasking Solution, Citrate-Based (Vector Lab Cat# H-3300-250), where the tissue is steamed for 40 mins. After cooling and washing in DI water, the slides were bleached to remove autofluorescence using a PBS-based solution containing 4.65% H2O2 and 35 mM NaOH and exposed to LED lights for 1 hr at room temperature. Following bleaching, slides were washed twice in 1x PBS and blocked in Intercept (PBS) Blocking Buffer (Li-Cor Cat# 927-70001) for 1 hr at room temperature.

Next, slides were incubated with unconjugated primary antibodies overnight (Panel 1, table), stained with fluorescently labeled secondary antibodies and Hoechst 33342 (1 µg/ml), coverslipped using 60% glycerol in 1X PBS, and then imaged using a CyteFinder (Rarecyte). After image acquisition, slides were placed in 1X PBS for at least 15 mins before de-coverslipping. To inactivate the fluorescence and allow for cycling, bleaching is also performed between cycles. Then, the slides were washed twice in 1x PBS and incubated with fluorophore-conjugated primary antibodies. The second serial section was used to stain with Panel 2 (Table).

For the serous intraepithelial carcinoma (STIC) samples, the tissue sections were first incubated in a 60°C oven for 3 hrs before continuing with dewaxing, rehydrating, and antigen retrieval steps. In addition, for the initial bleaching step before staining, the slides were bleached for 1 hr total but the PBS-based solution was refreshed at 30 mins. For all subsequent bleaching, the PBS-based solution was not refreshed midway through the step.

#### Image processing and quality control

Images collected on the CyteFinder were stitched and registered using the MCMICRO pipeline^90^ (https://github.com/labsyspharm/mcmicro). The UNMICST module^91^ in MCMICRO was used to segment images into single cells generating a data table containing mean fluorescence intensity for each marker in each cell. For the HGSOC samples, the regular UNMICST-solo was used, where a single DNA channel is used to determine cell nuclei. For the STIC samples, UNMICST-duo was chosen, using both a DNA channel and a nuclear envelope stain (lamin B1) to help differentiate between the tightly packed and irregularly shaped nuclei in STIC lesions. Accurate segmentation was assessed by visual inspection. Tissues were also visually inspected for cell loss during cycling, staining artifacts, and deformations. Antibodies that failed to stain or showed inaccurate staining patterns were excluded from downstream analyses.

### Preparing the Datasets

To generate an initial labeling of cell types in each dataset, SciMap’s rescale algorithm was used to standardize the signal intensity information of cell type markers, and then the phenotype_cells algorithm produced the initial labels. This labeling was then corrected with gates (binarized labels of expression / non-expression of a marker) and visual confirmation on respective CyCIF images. The corrections were done with a Boolean-based logic, in which a cell assigned an incorrect label would be assigned a different appropriate label or removed from the dataset based on combinations of gates. For non-cell type antibodies (e.g., ORF1P), markers were individually gated for each sample using Gator (https://github.com/labsyspharm/gater). Further correction to remove detected artifacts was then performed using a Latent Dirichlet Allocation (LDA)-based step. The raw signal intensity of all markers from all cells was used to generate a document-term matrix, and this matrix was used to generate latent weights which were then clustered to generate neighborhood labels for each cell. Cells in neighborhoods capturing artifact staining (as based on visual confirmation) were removed. The LDA procedure was implemented using the gensim package from Python, and clustering of the latent weights was done using the scikit-learn implementation of the k-means algorithm from Python.

Interferon signal intensity in each cell was measured and assigned a composite score based on the expression of the relevant markers (IFIT1, MX1) using a custom Python script. This script implemented the single-sample gene set enrichment analysis (SSGSEA) algorithm as defined by Barbie et al^92^. The scores outputted by the script were then binarized based on visual comparison against respective CyCIF images so that cells could be assigned labels of expression or non-expression (ISG^+^ or ISG^-^).

#### K-Means analysis

Clustering was performed with Python’s scikit-learn implementation of the k-means algorithm using the raw signal intensity of IFIT1 and MX1. Four clusters were made on each dataset; this optimal number of clusters was confirmed using elbow plots. Average raw signal intensity of IFIT1 and MX1 were calculated per cluster to relabel clusters based on ISG intensity.

In each sample, relative frequencies of cell types were counted per cluster. Then, for each cell type of interest, Wilcoxon’s signed rank tests were used to compare the normalized counts from the clusters with the highest and lowest levels of interferon signal intensity.

#### Spatial Counts

The spatial_count function from SciMap^41^ was used to measure the frequency of cell types and binarized interferon SSGSEA labels in the 20 µm radial neighborhood of each cell in each sample. First, tumor cells and CD8+ T cells were selected and any tumor cells or CD8+ T cells without neighbors in the query radius were excluded. Then, on a per-sample basis, for ISG+/- tumor cells and ISG+/- CD8+ T cells, the average frequencies of the neighbors’ cell types and interferon SSGSEA labels were calculated. Thus, this calculation generated a count of neighbors on the “representative” ISG+/- tumor cell and ISG+/- CD8+ T cell in each sample.

Wilcoxon’s signed-rank tests were used to statistically compare the counts of the ISG^+^ immune cell neighbors on the representative ISG^+^ and ISG^-^ tumor cells, and to compare the counts of the ISG^+^ tumor cell neighbors on the representative ISG^+^ and ISG^-^ CD8^+^ T cells.

#### Spatial Interactions

The spatial_interaction command from SciMap was utilized to measure the spatial association of ISG+/- tumor cells with each other on each sample. The query radius was set to 20 microns. This function generates randomized permutations of the interferon labeling on the tumor cells in each sample and then calculates the probability of adjacency for the two classes of tumor cells^41^.

### Statistical Tests

The scipy package in Python (v3.10.9) was used to perform Pearson’s R correlations (pearsonr function) and Wilcoxon’s signed-rank tests (wilcoxon function). For the comparison of ISG^+^ CD8 T cell proportions in ISG^high^ compared to Negative k-means clusters, we used a Mann-Whitney test in GraphPad (10.0.2).

## Acknowledgements

This work was funded by the Dr. Miriam and Sheldon G. Adelson Medical Research Foundation (JSB, BJNW, RD, DJS, GEK), the Ludwig Center at Harvard (JSB), the Ovarian Cancer Research Alliance Liz Tilberis Early Career Awards (BJNW), the Ovarian Cancer SPORE P50 CA228991-06A1 (RD), and a University of Minnesota Grand Challenges Grant (BJNW). Research at the University of Minnesota was supported, in part, by the National Institutes of Health’s National Center for Advancing Translational Sciences, grant UM1TR004405 and the Masonic Cancer Center’s shared resources grant NIH P30 CA77598, the University of MN Grand Challenges Grant (TKS, BJNW, MS, and ACN), the Jan Chorzempa Cancer Research Endowed Fund (TKS) and the University of MN Masonic Cancer Center Translational Working Group Grant (TKS, BJHW). From HMS, we thank Laura Selfors and Francesca Silvestri for bioinformatics support, Peter Sorger and Sandro Santagata for advice and support on imaging studies, other members of the Laboratory for Systems Pharmacology for spatial imaging advice and support, including Jia (Jerry) Lin, Edward Novikov, Peter Barera, and Jeremy Muhlich for valuable feedback for image analysis and O2 support, Zoltan Maliga for antibody validation, and Tanjina Kader advice on STIC imaging. We also thank Pamoda Galhenge and Enakshi Sunasse for helpful discussions, Jeff Yu for technical assistance, and Brendan Shay and Grace Gao for lab management. From UMN, we thank Marissa Macchietto and Christy Henzler (University of Minnesota Super Computing Institute) for computational/statistical advice, Emily Stock for assistance with patient record analysis.

## Extended Data

**Extended data figure 1.**
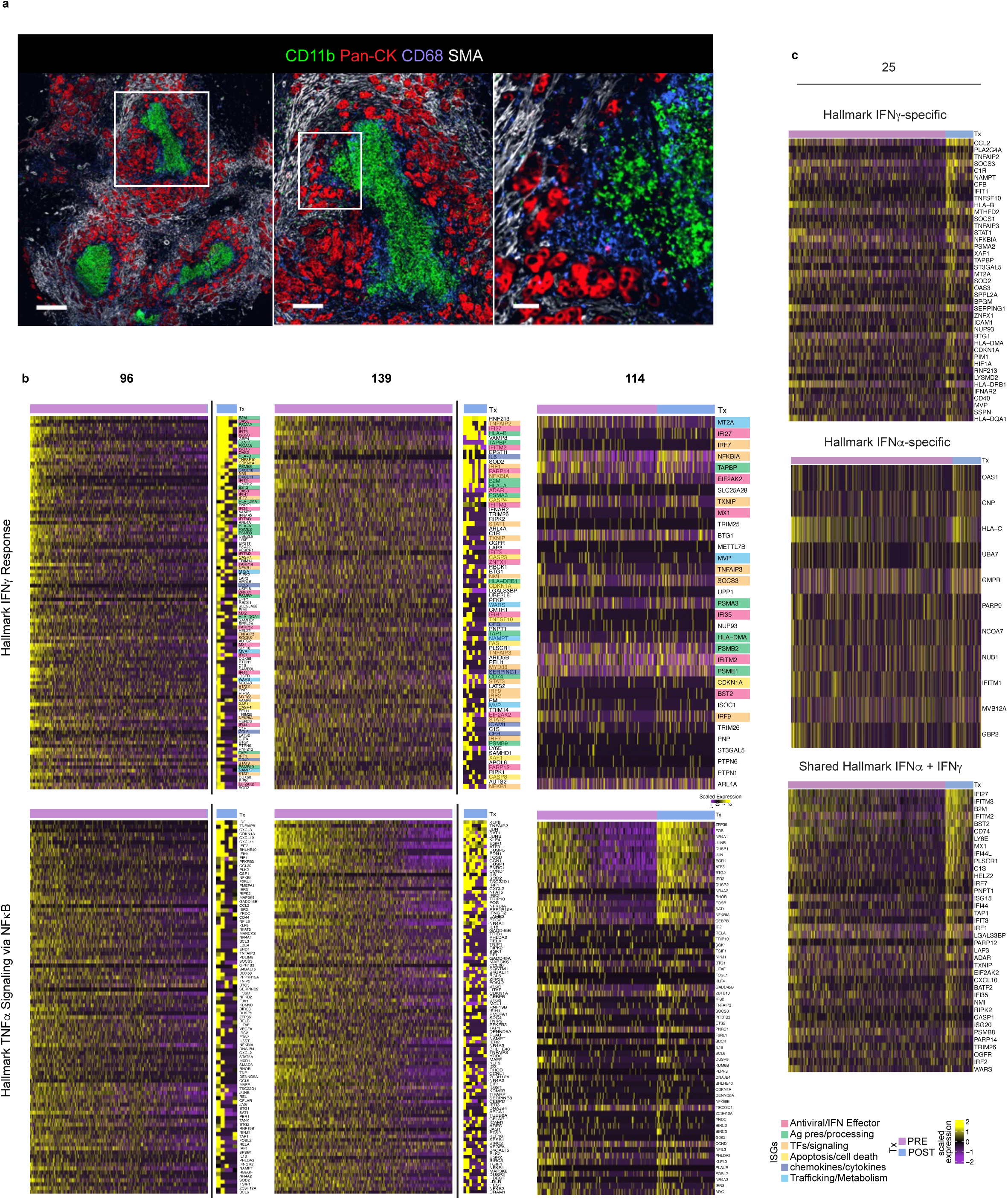
Analysis of IFN and TNFA/NFKB signatures and spatial imaging of tumor 38. a. Cyclic immunofluorescent (CyCIF) images of patient 38 pre-treatment tumor showing the myeloid clusters (CD11b^+^, green) surrounded by tumor cells (panCK^+^, red) with abundant surrounding stroma (SMA^+^, white). Left panel showing multiple myeloid clusters (scale bar = 500 µm). Middle panel showing enlargement of white box in left panel (scale bar = 200 µm). Right panel showing enlargement of white box in middle panel (scale bar = 50 µm). b. Heatmaps showing the level of expression of differentially expressed Hallmark IFNγ and ‘TNFα signaling via NFκB signature’ genes in the three paired samples with the low post-treatment cell counts (patients 96 and 139) or low upregulation of IFNγ in post-treatment cells (patient 114). Cells were clustered by pre- and post-treatment (purple and blue bars at top). Genes were rank-ordered in the maps based on the cluster average fold changes between post- and pre-treated tumors. Color coding represents the different classes of IFN signature genes as in Figure 1 (see legend). The scale for the post treatment cells in patient 139 and 96 tumors are enlarged in order to visualize the small numbers of cells. c. Heatmaps showing genes that are specifically associated with the Hallmark IFNγ signature (top panel), the IFNα signature (middle panel), or those common to both (bottom panel) in patient 25.

**Extended data figure 2.**
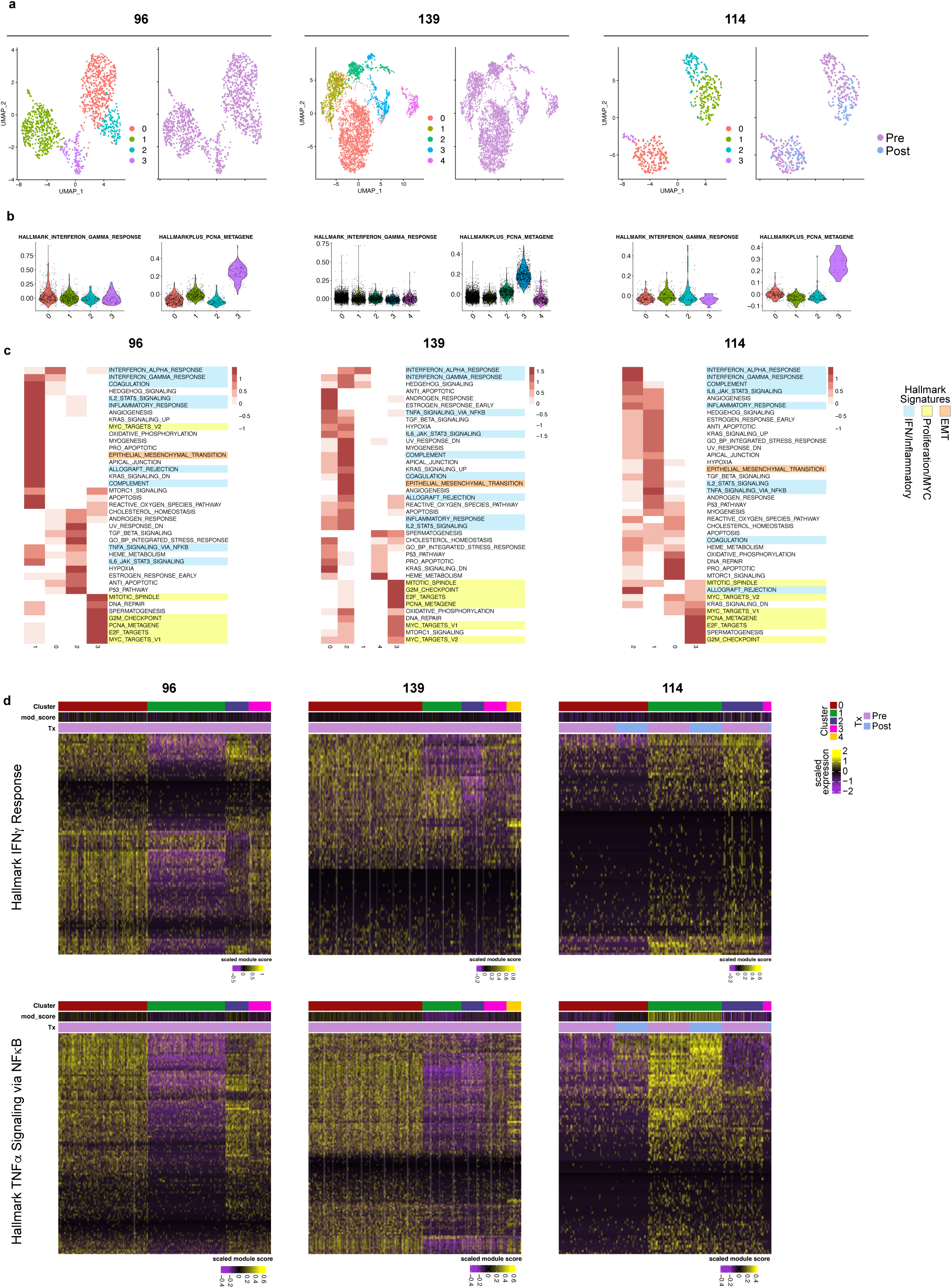
Analysis of cancer cell clusters in additional pre-post tumor pairs. a. UMAP plots of cancer cell clusters in three pre-post tumor pairs. Colors denote the cluster numbers (upper) or the pre-post status of each of the cells (lower). b. Violin plots of the IFNγ and PCNA signature scores for each cluster. c. Heatmaps showing the signature scores for the pseudo-bulk average gene expression of each cluster for 39 gene sets. Gene sets are colored by functional category (blue: IFN/IFM genesets, yellow: proliferation-related genesets, and orange: EMT). d. Heatmaps showing the expression of the Hallmark IFNγ response or TNFA/NFKB signature genes in each cluster (as in Figure 3).

**Extended data figure 3.**
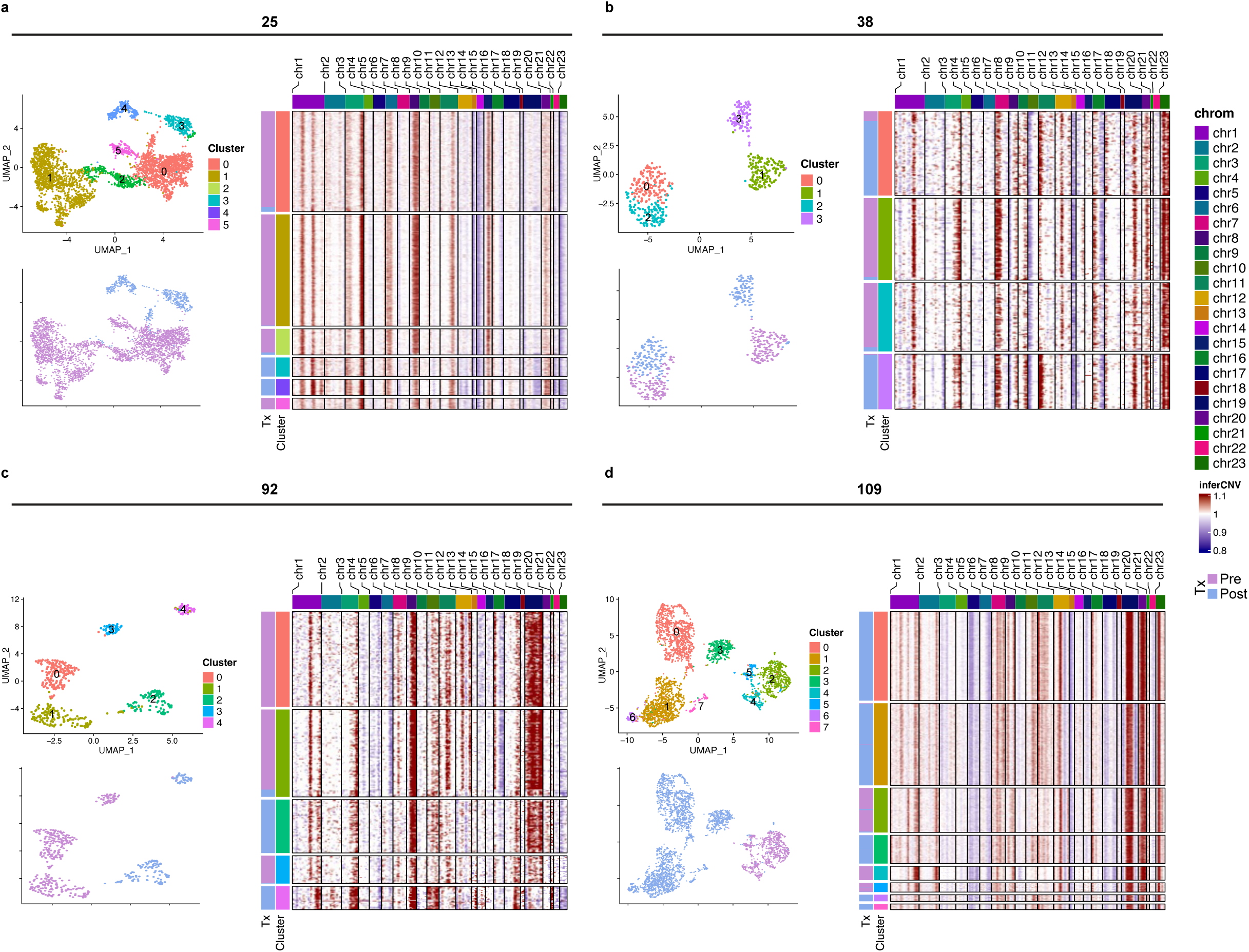
Analysis of changes in copy number variation in cancer cells from patients before and after therapy. UMAP plots colored by cluster (top left) or by pre- vs post-treatment (bottom left) and inferred CNAs for individual cells (right panel) for patients 25 (a), 38 (b), 92 (c), and 109 (d). Graphic of CNAs has cells (rows) arranged by cluster, with sidebar indicating which cells are pre- vs post-therapy (Tx). Copy number gains are in red and losses are in blue (see legend).

**Extended data figure 4:**
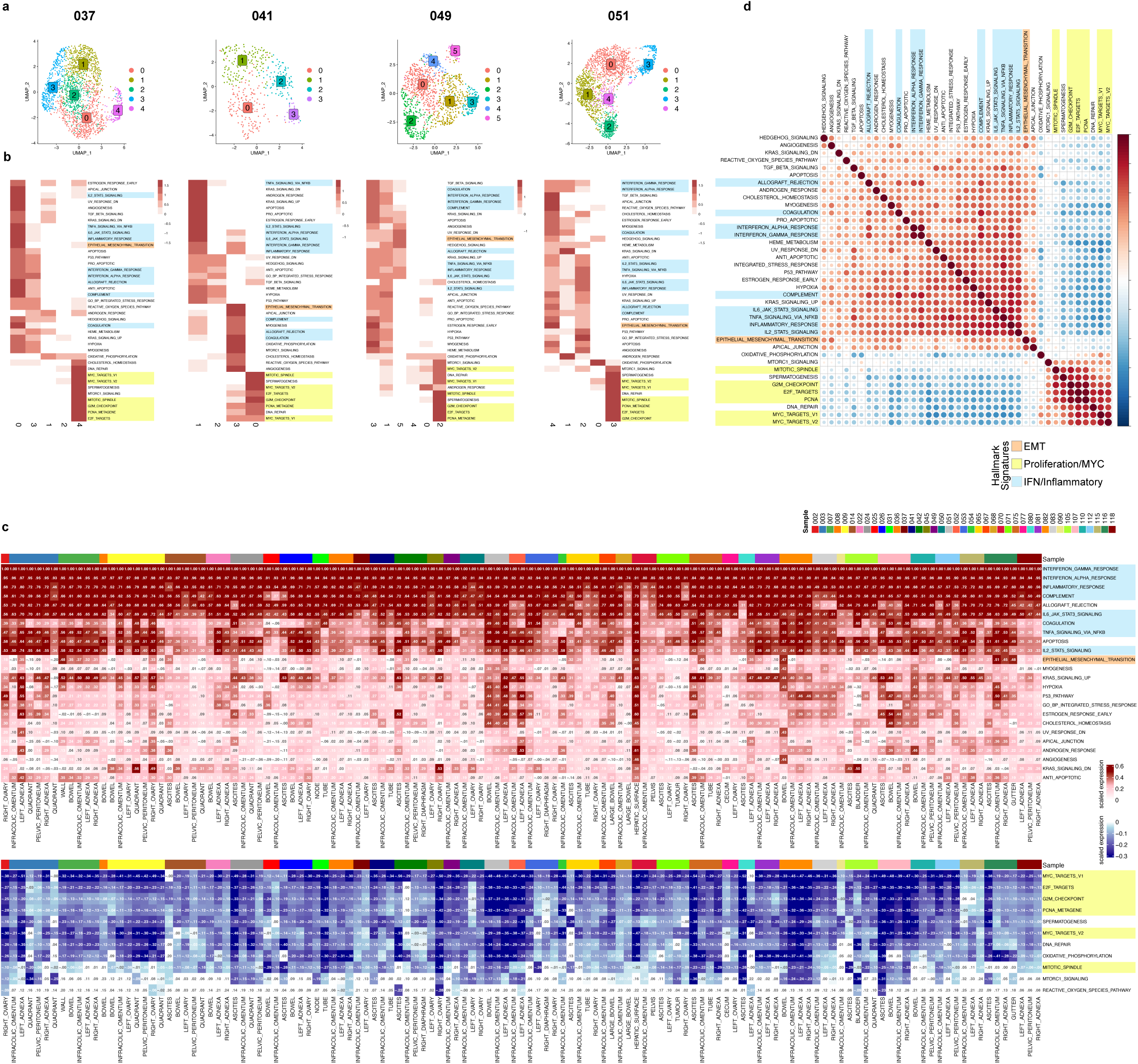
Analysis of Pre-treatment tumors from the MSKCC SPECTRUM dataset. a. UMAP plots of single epithelial cell clusters derived from four representative pre-treatment tumor samples from omental sites, color-coded by cluster. b. Heatmaps showing the signature scores for the pseudo-bulk average gene expression of each cluster. Gene sets are colored by functional category as in Figure 2. c. Table showing the Pearson correlation value for each signature compared to the IFNγ signature. Colored bar designated the patient number for each of the metastatic sites examined. d. Matrix showing the pairwise Pearson correlation values based on GSEA scores.

**Extended data figure 5.**
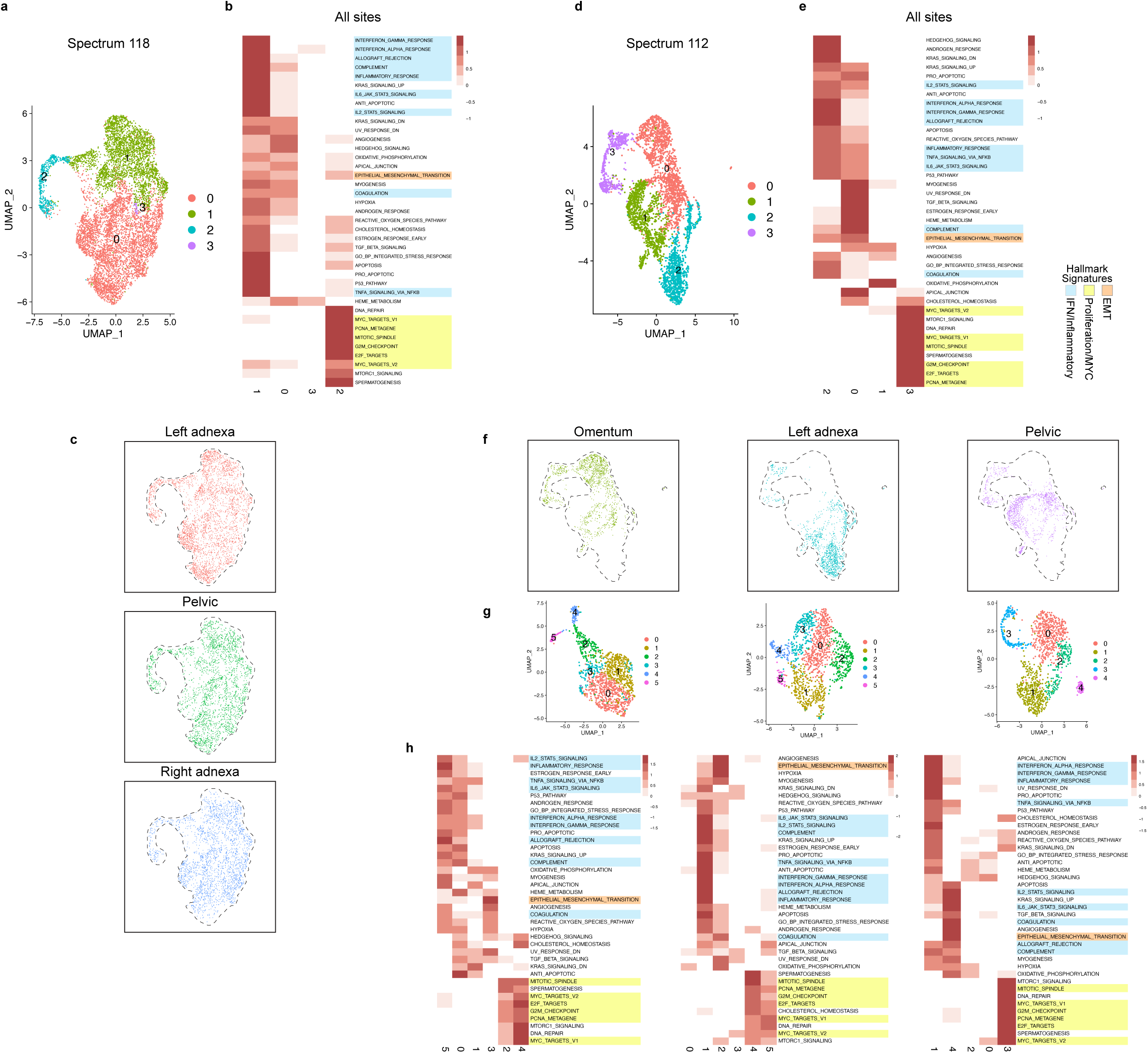
Analysis of tumor clusters at different metastatic sites in the MSK SPECTRUM dataset. a. UMAP of integrated and clustered scRNAseq data from all metastatic sites from patient 118. b. Heatmap of scaled signature scores for the pseudo-bulk average gene expression of each cluster in (a) for 39 gene signatures. c. UMAP from (a) colored by cells from each metastatic site. d. UMAP and e. heatmap for patient 112 generated as in (a-b). f. UMAP from (d) colored by cells from each metastatic site. g. UMAPs of clusters generated from each metastatic site separately. h. Heatmaps of scaled signature scores for the pseudo-bulk average gene expression of each cluster in (g). Gene signatures are colored by functional category as previously described.

**Extended data figure 6.**
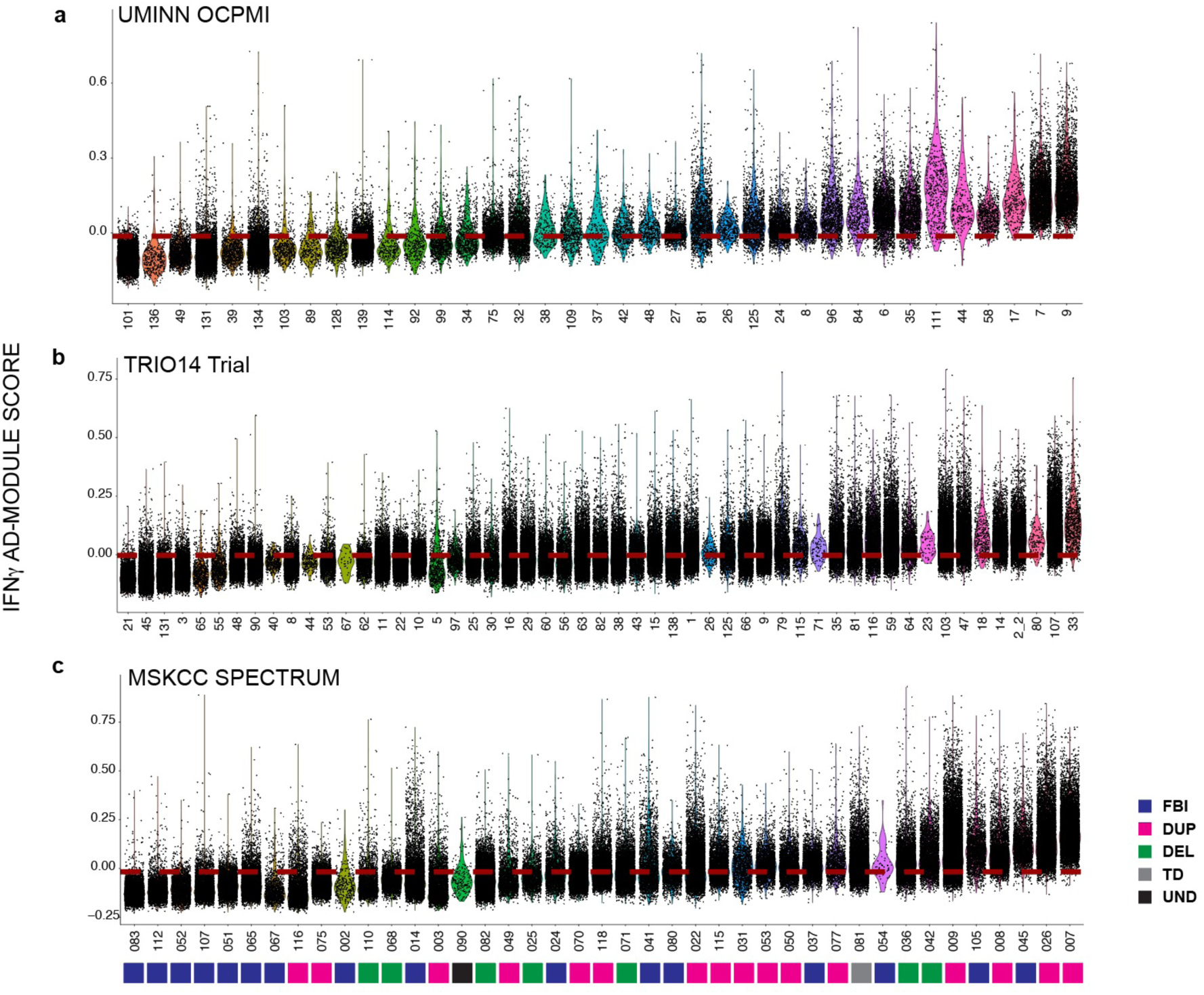
IFNγ signature scores from each patient in the UMINN-OCPMI, TRIO14, and MSKCC SPECTRUM datasets. IFNγ signature scores for each tumor in the UMINN OCPMI (a), TRIO14 (b) and MSKCC SPECTRUM (c) datasets were determined and plotted in the volcano plots shown. For the SPECTRUM dataset samples, all metastatic sites were included for each patient sample. The samples are ordered by the number of cells above median IFNγ value (shown in red dashed line). For the SPECTRUM dataset, the bars below each sample indicate the type of mutational signature associated with the tumor as previously reported^50^. FBI (fold back inversion bearing tumors), HRD-DUP (homologous recombination defective *BRCA1* mutant-like), HRD-DEL (homologous recombination defective *BRCA2* mutant-like), TD (tandem duplication), UND (undetermined).

**Extended data figure 7.**
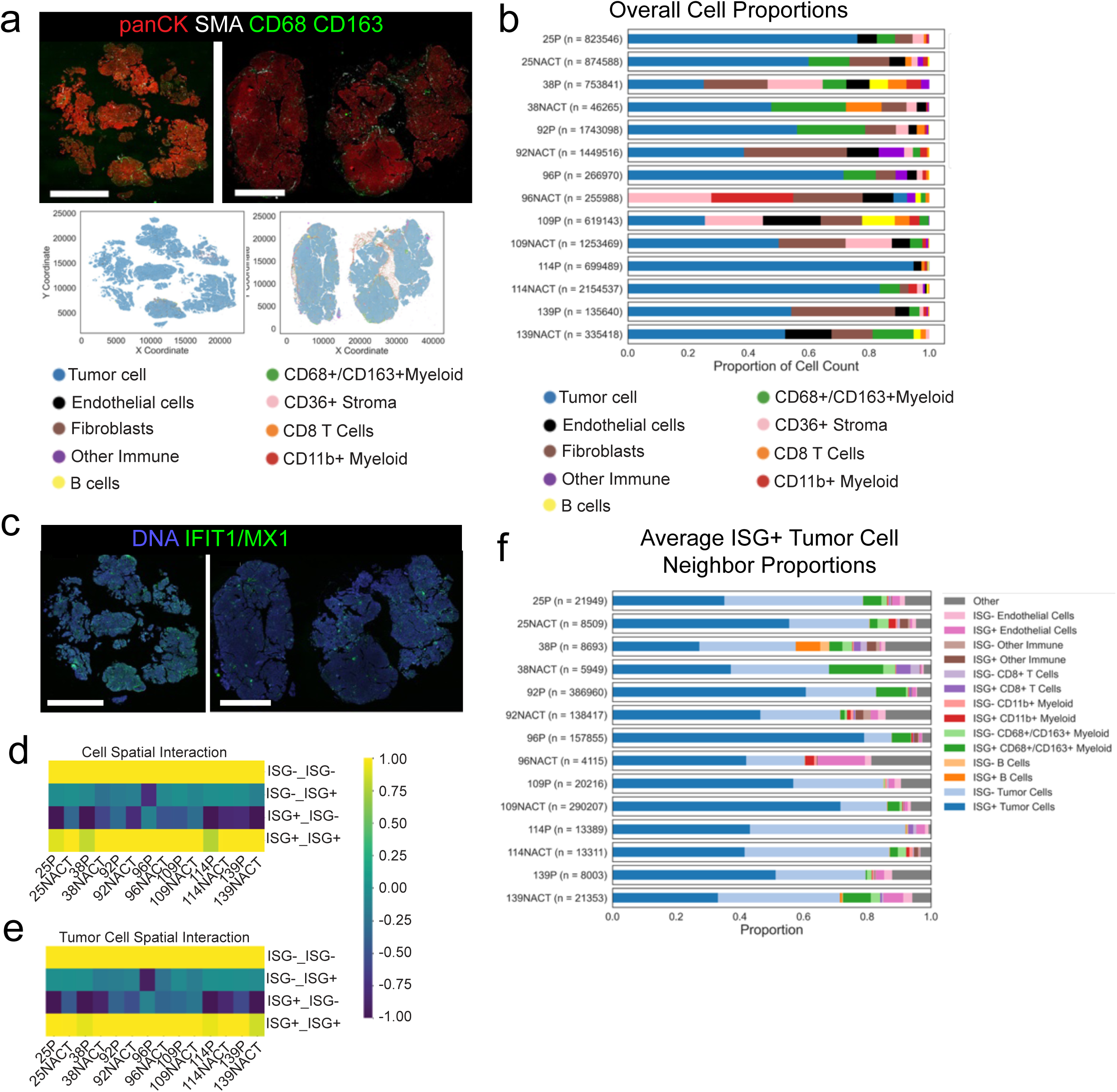
CyCIF analysis of patient tumors. a. CyCIF images (top; red=panCK, white=SMA, green=CD68/CD163) and cell type centroid plots (bottom) are shown for pre-(left) and post-treatment (right) tissue from patient 114. Scale bars = 5 mm. b. The quantification of cell types for all patients is shown. The number in parentheses indicates the number of cells analyzed for that tissue. c. CyCIF images showing IFIT1 and MX1 (green) staining in pre-(left) and post-treatment (right) tissue from patient 114. Scale bars = 5 mm. d. Heatmap showing cell association based on ISG status, irrespective of cell identity. e. Heatmap showing tumor cell association based on ISG status. f. The average proportions of ISG^+^ tumor cell neighbors within a 20 µm radius for all patients are shown in a stacked bar plot. Total queried cells are shown in parentheses.

**Extended data figure 8.**
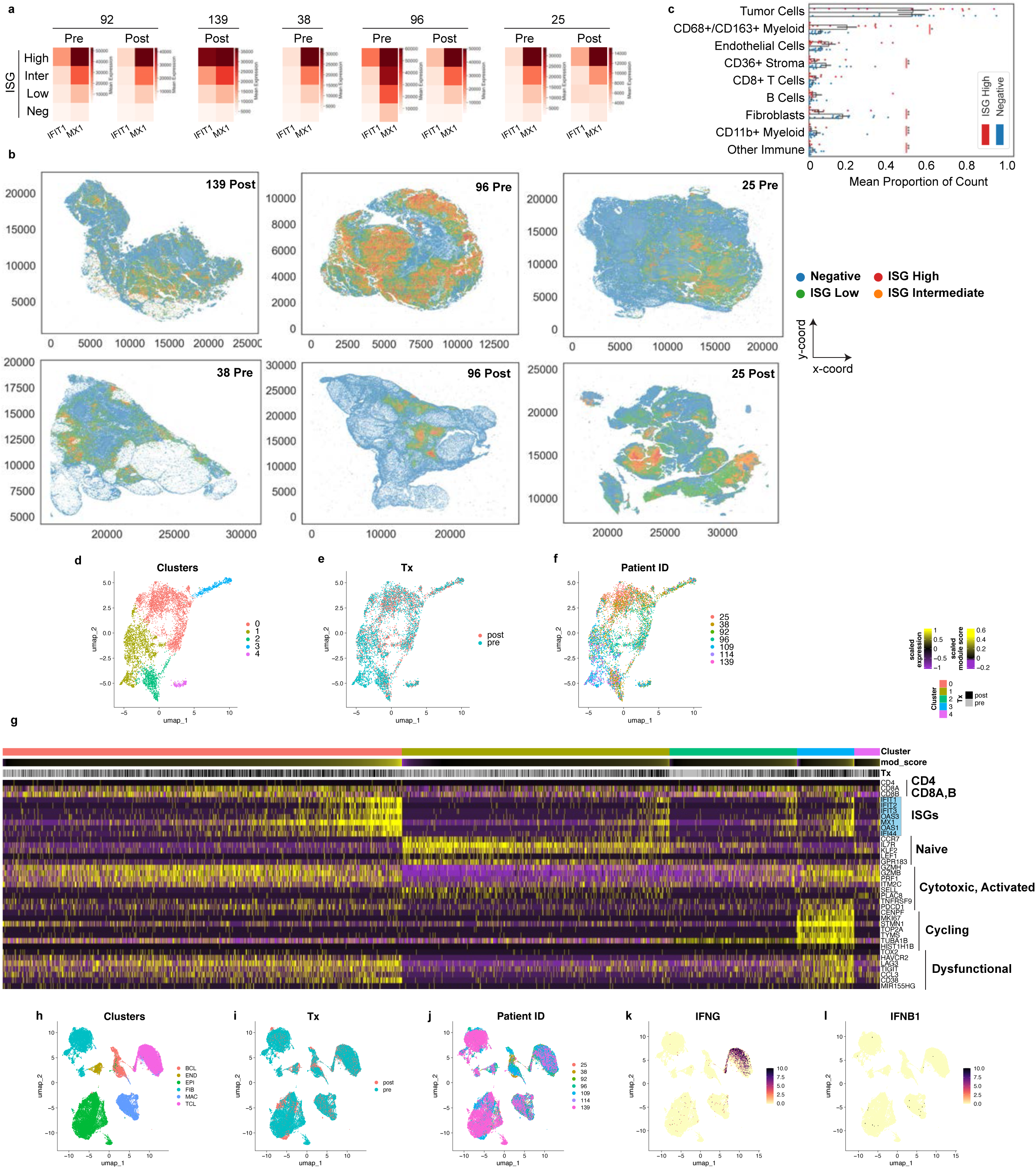
Analysis of ISG zones in patient tumors and analysis of CD8^+^ T cell subtypes that express ISGs. a. Heatmaps are presented for the indicated tissues showing cell clusters based on IFIT1 and MX1 intensities. b. Centroid plots show the spatial distribution of k-means clusters found in (a). c. The quantification of cell type proportions in the ISG^high^ and the Negative k-means clusters. Wilcoxon signed rank tests were used to compare the normalized counts. p<0.05 *, p<0.01 **, and p<0.001 ***. UMAPs of CD8^+^ T cells from all patients by cluster (d), treatment status (e), and individual patients (f). g. Heatmap of CD8^+^ T cells including gene markers of different CD8 T cell subtypes. Cells are binned in clusters as in d (top bar), rank ordered by IFNγ score (2nd bar), and indicated as pre (grey) or post (black) (3rd bar). UMAPs of all cell types from all patients by cluster (h), treatment status (i), and individual patients (j). UMAP visualizing IFNG (k) and IFNB1(l) expression.

**Extended data figure 9.**
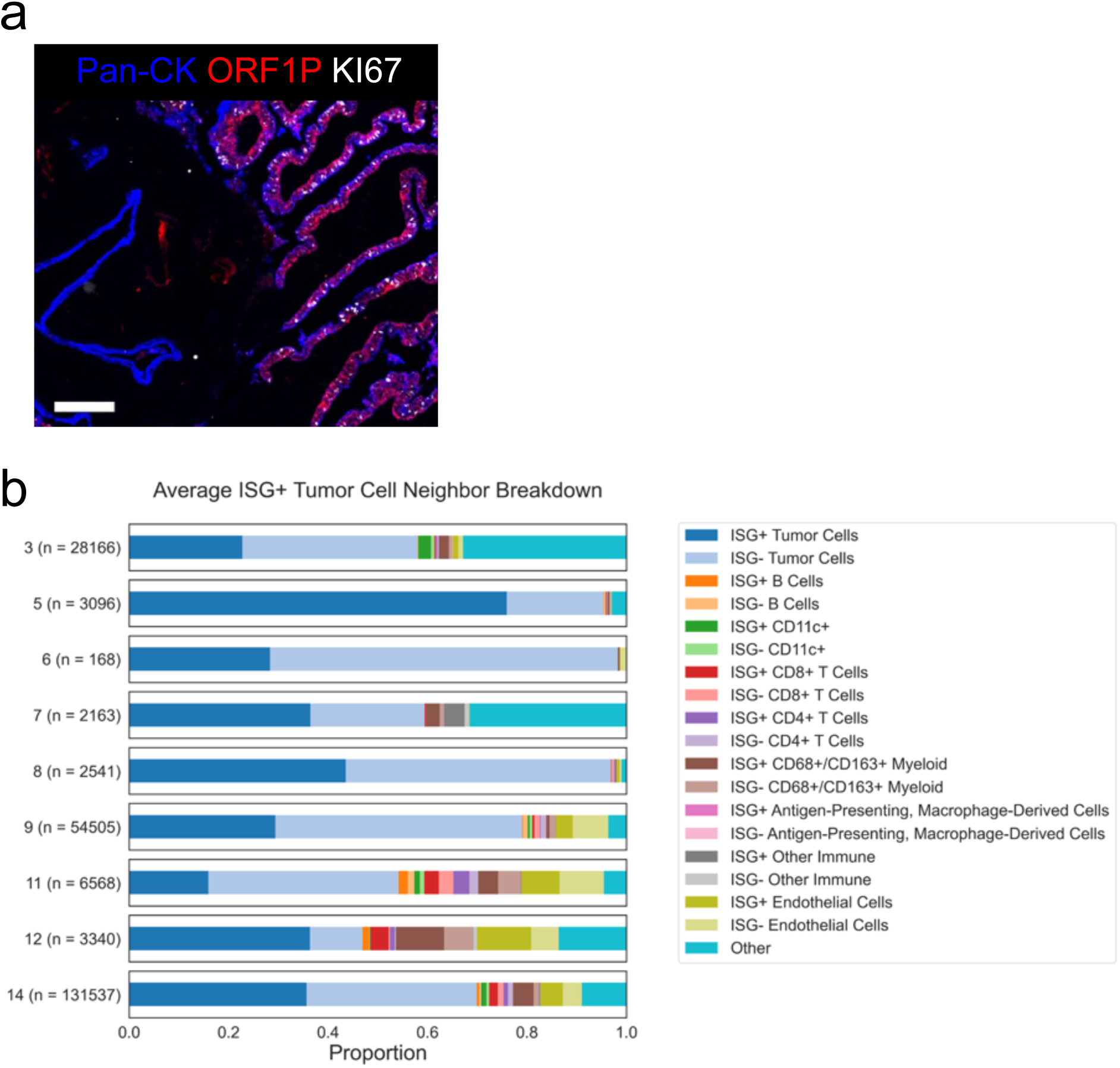
Analysis of ORF1P in non-STIC regions of a fallopian tube and neighbor analysis of ORF1P^+^ tumor cells in fallopian tubes. a. CyCIF image of region with normal fallopian tube epithelium and STIC lesions. PanCK (blue) marks epithelial cells and Ki67 (white) shows high proliferation in STIC lesion. The ORF1P (red) signal is restricted to the STIC. Scale bar = 200 µm. b. The average proportions of ISG^+^ tumor cell neighbors within a 20 µm radius for all patients whose fallopian tube tissue included a tumor are shown in a stacked bar plot. Darker shades of the same color indicate ISG^+^ cells within a cell type. Total queried ISG^+^ tumor cells are shown in parentheses.

## Supplemental Information

**Supplemental data figure 1. Lack of an association between IFN and DNA damage markers and Jaccard similarity coefficients between gene sets.** a. CyCIF image showing tumor cells (pan-CK^+^, blue) that either express ISG IFIT1 (green, overlay cyan) or do not. In these areas, phospho-γH2AX positivity (red) was monitored to assess co-expression of IFIT1 and γH2AX. Arrowheads denote yH2AX-positive cells. b. Pearson correlation values of IFIT1 vs γH2AX and MX1 vs γH2AX are presented for all of the matched tissues. c. CyCIF images showing IFIT1 (green) and the nuclear marker, BAF (red), which when punctate indicates micronuclei. The image was counterstained with Hoechst (blue). White dashed lines show the interface between IFIT1^+^ areas and BAF dense areas. In all CyCIF images, scale bar = 50 µm. Patient samples are indicated on corresponding images. d. Jaccard similarity coefficients (from 0 to 1) between 50 Hallmark gene sets. Pairs of gene sets with correlations > 0.10 are highlighted with black boxes.

**Supplemental data figure 2: Analysis of tumors from the Trio14 trial.** a. UMAPs of single epithelial cell clusters derived from pre-treatment tumor samples color-coded by cluster. b. Heatmaps showing the signature scores of pseudo-bulk average expression of each cluster. c. Table showing the Pearson correlation for each signature compared to the IFN*γ* signature in each sample. d. Matrix showing the pairwise Pearson correlation values based on GSEA scores. Gene sets are colored by functional category as described in Figure 2.

**Supplemental data figure 3. Analysis of expression of IFN**γ **and Inflammatory Response gene signatures in B cells.** Violin plots and gene expression heatmaps of the Hallmark IFNγ and Inflammatory response gene sets for patients 25, 38, 92 and 109 in B cells. Cells are grouped by pre-(purple) vs post-(blue) treatment in both the violin plots and the heatmaps and ordered by highest to lowest IFNγ score in each group in the heatmaps. Samples 92 and 109 contained either none or a few post-treatment cells.

**Supplemental data figure 4. Analysis of expression of IFN**γ **and Inflammatory response gene signatures in patient tumor 96.** Violin plots (a, c) and gene expression heatmaps of the Hallmark IFNγ (b) and Inflammatory response (d) gene sets for patient 96 in tumor cells, T cells, myeloid cells, fibroblasts and endothelial cells. Cells are grouped by pre-(purple) vs post-(blue) treatment in both the violin plots and the heatmaps and ordered by highest to lowest IFNγ score in each group in the heatmaps.

## Supplementary Tables

Supplementary Table 1: Differentially expressed genes (DEGs) for each matched pair

Supplementary Table 2: Gene set enrichment analysis (GSEA) for each matched pair

Supplementary Table 3: Most commonly enriched genes in the different Hallmark signatures

Supplementary Table 4: Hallmark IFNγ_Response and TNFα_Signaling_Via_NFκB in tumor cells

Supplementary Table 5: Hallmark and Custom Gene Sets

Supplementary Table 6: Hallmark IFNγ_Response and Inflammatory_Response in stromal cells

Supplementary Table 7: CyCIF antibody panels

