## Supplementary figures and images for "Tumor Cells Enriched for Interferon and Inflammatory Programs Pre-Exist in High Grade Serous Ovarian Cancer and are Proportionately Significantly Increased Post Chemotherapy"

### Supplemental figure 1

##### Supplemental Figure 1

**a**

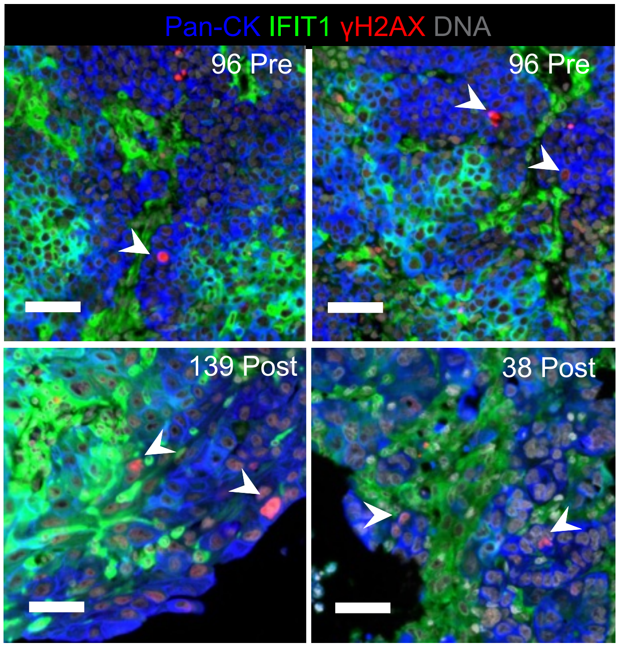


**c**

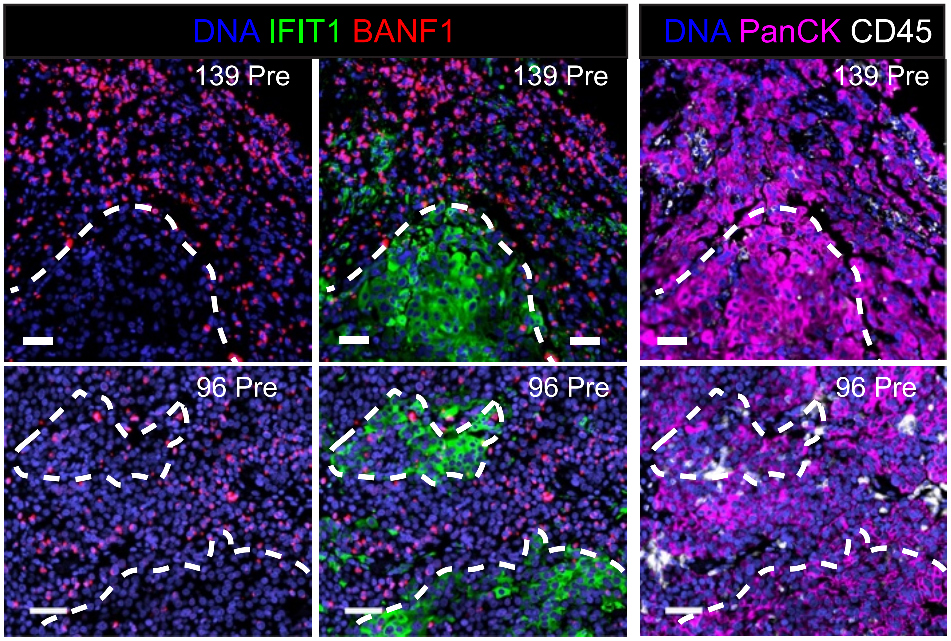


**b**

| | IFIT1 vs H2AX | MX1 vs H2AX |
|---|---|---|
| 25P | 0.10 | 0.03 |
| 25NACT | -0.01 | 0.03 |
| 38P | 0.03 | -0.02 |
| 38NACT | 0.03 | 0.03 |
| 92P | 0.03 | 0.01 |
| 92NACT | 0.04 | -0.01 |
| 96P | -0.07 | -0.13 |
| 96NACT | 0.03 | 0.00 |
| 109P | 0.03 | -0.04 |
| 109NACT | 0.01 | -0.02 |
| 114P | 0.22 | 0.10 |
| 114NACT | 0.13 | 0.10 |
| 139P | -0.02 | -0.02 |
| 139NACT | -0.03 | -0.05 |

**d**

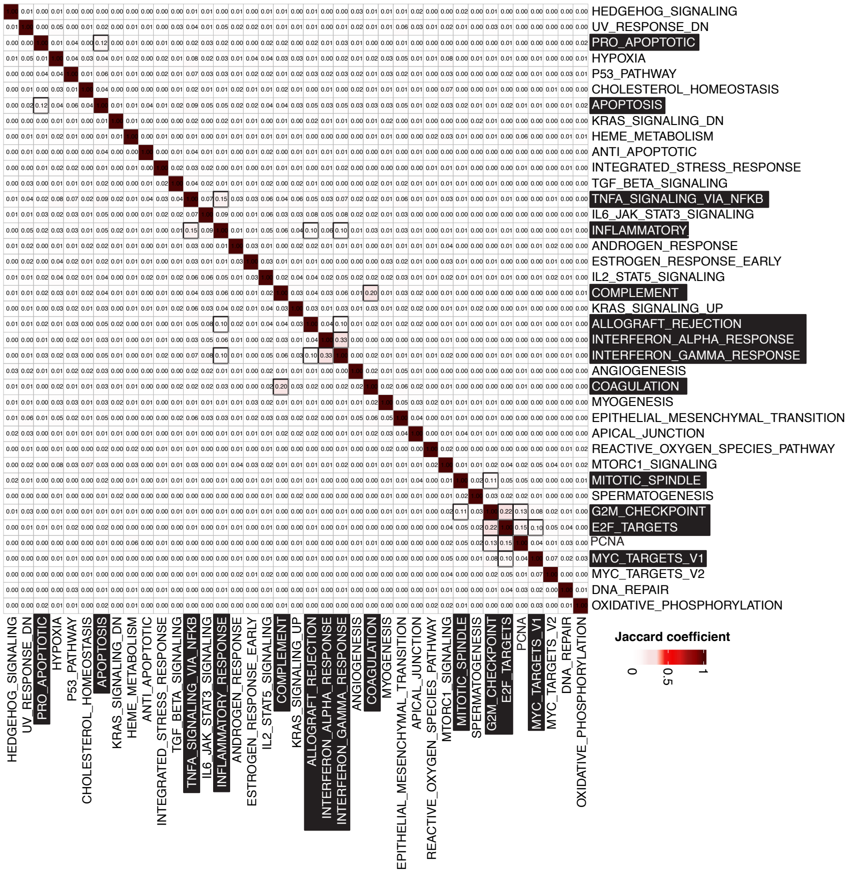

### supplemental figure 2

#### Supplemental data figure 2

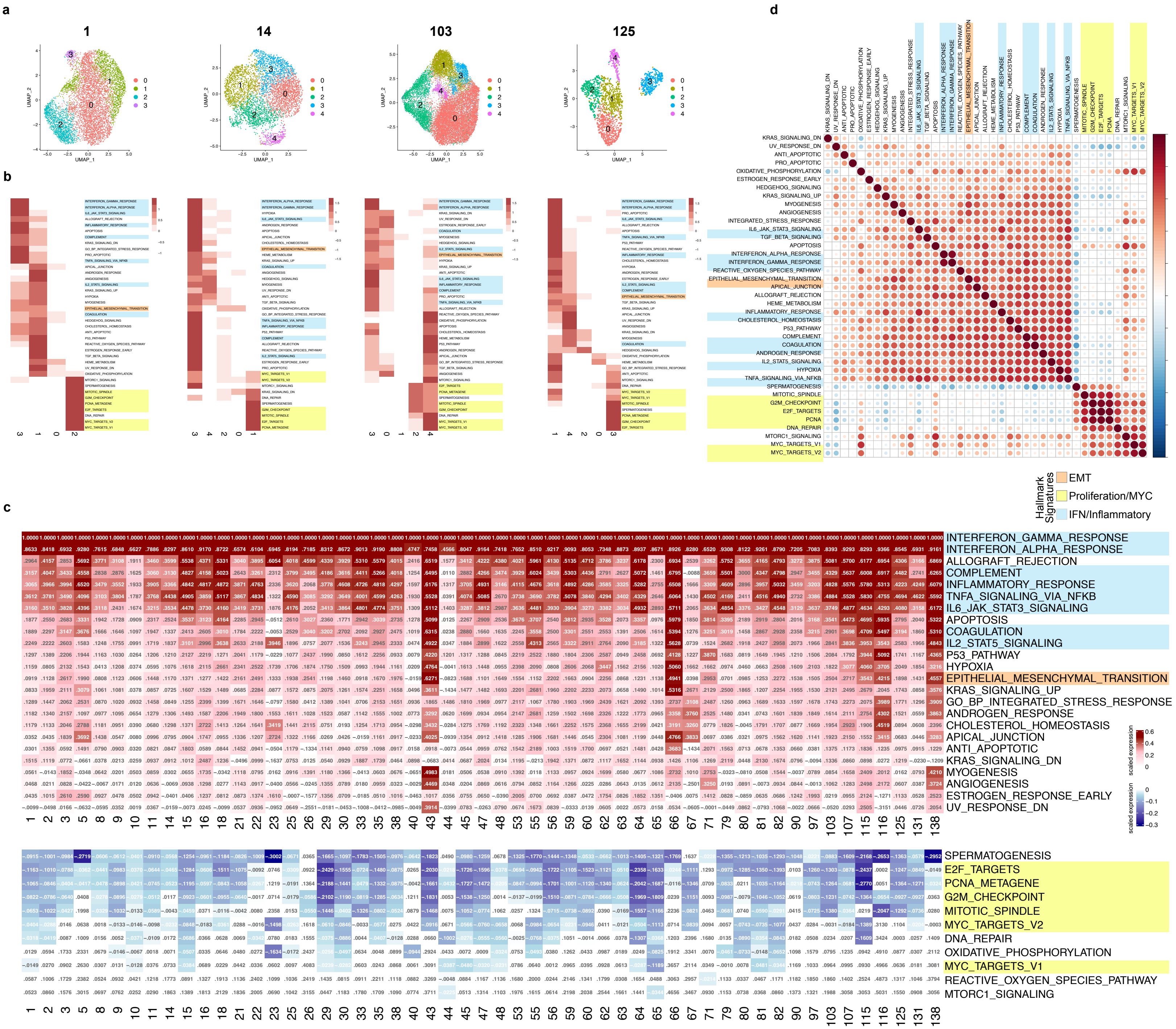

### supplemental figure 3

#### Supplemental data figure 3

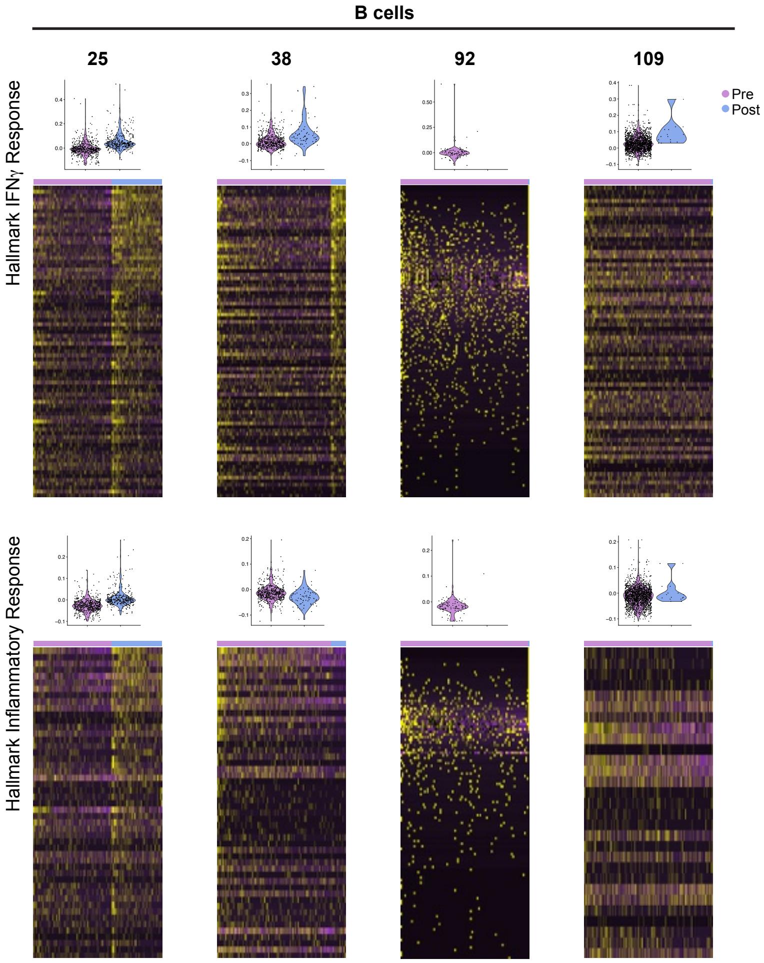

### supplemental figure 4

#### Supplemental figure 4

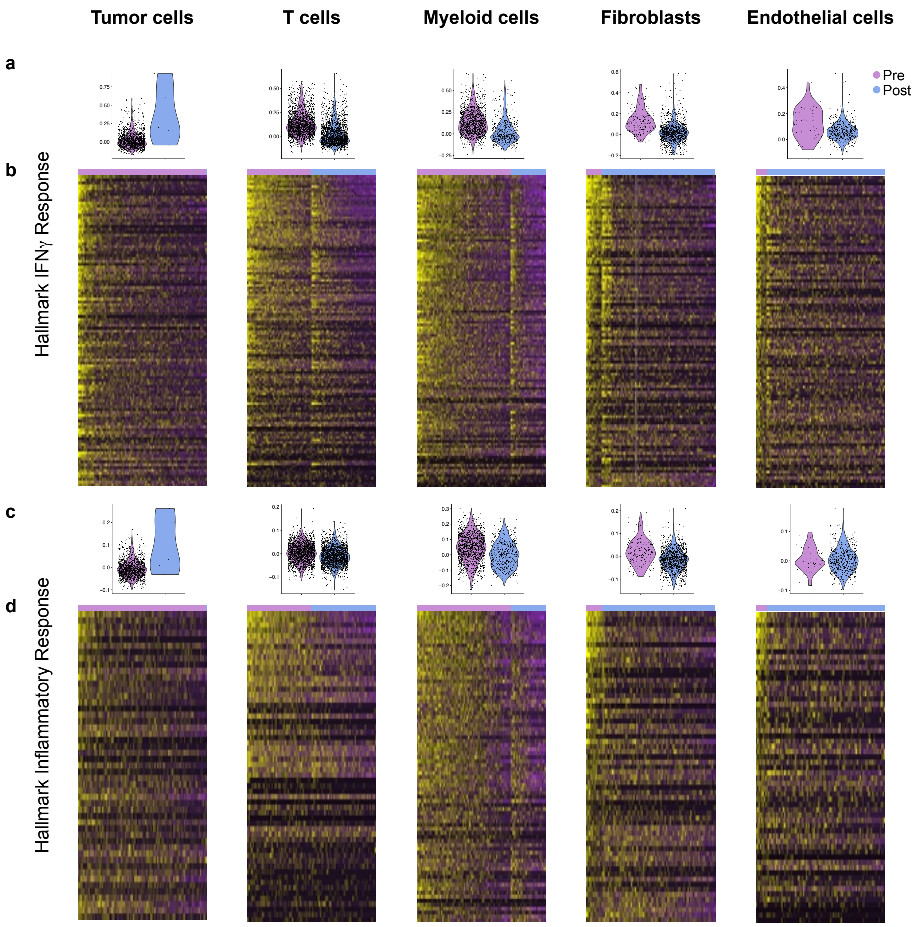
